# Chromatin assembly by the histone chaperone HIRA facilitates Human Papillomavirus replication

**DOI:** 10.1101/2025.09.05.674520

**Authors:** Ashley N. Della Fera, Dan Chen, Alison A. McBride

**Affiliations:** Laboratory of Viral Diseases, National Institute of Allergy and Infectious Diseases, 33 North Drive, MSC3209, National Institutes of Health, Bethesda, Maryland 20892, USA

**Keywords:** Keyword: HIRA, Histone variant H3.3, keratinocyte, HPV, human papillomavirus, viral replication, transcription, epigenetics, H3.3 phosphorylation

## Abstract

The circular, double-stranded DNA genomes of Human papillomaviruses (HPV) exist in a nucleosomal state throughout the infectious cycle and rely on host histone epigenetic modifications and chromatin assembly processes to promote various phases of the viral life cycle. Here, we show that the histone H3.3 chaperone HIRA and its associated complex members are recruited to HPV replication factories during the late phase of the HPV life cycle. HIRA is also recruited to HPV replication factories generated by amplification of a replicon with a minimal origin and expression of the viral replication proteins E1 and E2, demonstrating that the E1 and E2 proteins are sufficient for HIRA recruitment. Downregulation of HIRA expression reduces HPV31 DNA amplification and viral transcription in differentiated keratinocytes. Histone H3.3 that is highly phosphorylated on serine residue 31 is also enriched at sites of HPV replication and this modification links the DNA damage response to chromatin that supports rapid gene activation. We propose that deposition of histone H3.3 generates viral minichromosomes that are highly primed to support the late stages of the HPV life cycle.

## 1. Introduction

Human papillomavirus minichromosomes are assembled in chromatin at all stages of the viral infectious cycle. Assembly of nucleosomes onto DNA is a controlled process facilitated by replication-dependent and replication-independent nucleosome assembly pathways. Histone chaperones function in these processes to deposit either canonical histones or histone variants during DNA replication, repair and active gene transcription, or to aid in the maintenance of heterochromatin DNA [1]. Histone H3.1 is primarily deposited on chromatin during DNA replication in S-phase while H3.3 is deposited in a replication-independent manner. Replication-independent histone H3.3 deposition is facilitated by two histone chaperone complexes, Daxx/ATRX and HIRA/UBN1/CABIN1 [2]. The Daxx/ATRX complex aids in the maintenance of heterochromatin DNA at telomeres and peri-centromeres, while the HIRA/UBN1/CABIN1 complex, in cooperation with the histone H3.3/H4 binding protein ASF1a, functions in active processes such as DNA replication, DNA repair, and transcription [3].

Previous studies from our laboratory demonstrated that HPV genomes packaged in virions are enriched in the histone H3.3 variant relative to host chromatin [4]. This led us to question the role of the histone H3.3 chaperone HIRA in the HPV infectious cycle. HPVs amplify their genomes in differentiated cells that have exited the cell cycle and are in a pseudo G2-like phase [5, 6]. Thus, HPV genomes are dependent on replication-independent histone chaperones to assemble viral minichromosomes. Replication of HPV genomes during the late phase of the viral life cycle also utilizes host DNA repair machinery [7]. HIRA functions both locally and globally in DNA repair; HIRA deposits H3.3 into damaged chromatin [8] and promotes transcriptional recovery after DNA repair independently of deposition of H3.3 [9]. HIRA also deposits H3.3 onto transcriptionally active chromatin throughout the cell cycle and has been implicated in various pro-and anti-viral responses during infection with other DNA viruses. HIRA deposits histones on naked DNA [10] and can silence incoming HSV genomes [11], but conversely promotes Hepatitis B infection [12, 13]. HIRA is also implicated in the host innate immune response and, on occasion, can be found associated with promyelocytic leukemia nuclear bodies (PML-NBs) [14–17]. By contrast, Daxx is constituently associated with PML-NBs where it maintains a soluble pool of H3.3/H4 dimers and aids in the organization of heterochromatin [2].

Here, we show that HIRA is a positive regulator of late HPV replication and transcription. Moreover, we show that H3.3 is phosphorylated on serine 31 during the process of HPV replication and propose how this might promote high levels of late viral transcription.

## 2. Results

### 2.1 HIRA and Daxx associate with HPV16, HPV18, and HPV31 viral replication foci

To determine whether histone H3.3 chaperones play a role in late HPV DNA replication, we examined the localization of HIRA and Daxx in keratinocytes containing extrachromosomal HPV31 genomes. CIN612-9E cells were differentiated to induce the late phase of infection using a method described by Moody and colleagues [18]. The association of HIRA and Daxx with HPV replication foci was evaluated by immunofluorescence in differentiated CIN612-9E cells that amplify the HPV31 genome in replication factories (**Figure 1A-B**), and in differentiated human foreskin keratinocytes (HFKs) immortalized with and containing extrachromosomal HPV16, HPV18, or HPV31 genomes (**Figure 1C-E**). The single-stranded DNA binding replication protein (RPA2) was used as a marker for viral replication foci [19]. We observed localization of both HIRA and Daxx at >90% sites of HPV31 replication in differentiated 9E cells, with an average enrichment score of 1.9 and 6.2, respectively, compared to the surrounding nucleus (**Figure 1A** and **1B**). Moreover, both HIRA and Daxx localized at >88% HPV replication factories in differentiated HFK cells immortalized by HPV16, HPV18 and HPV31, with an average enrichment score of 1.8 and 4.4, respectively, as compared to the nucleoplasm (**Figure 1C-E**).

**Figure 1:**
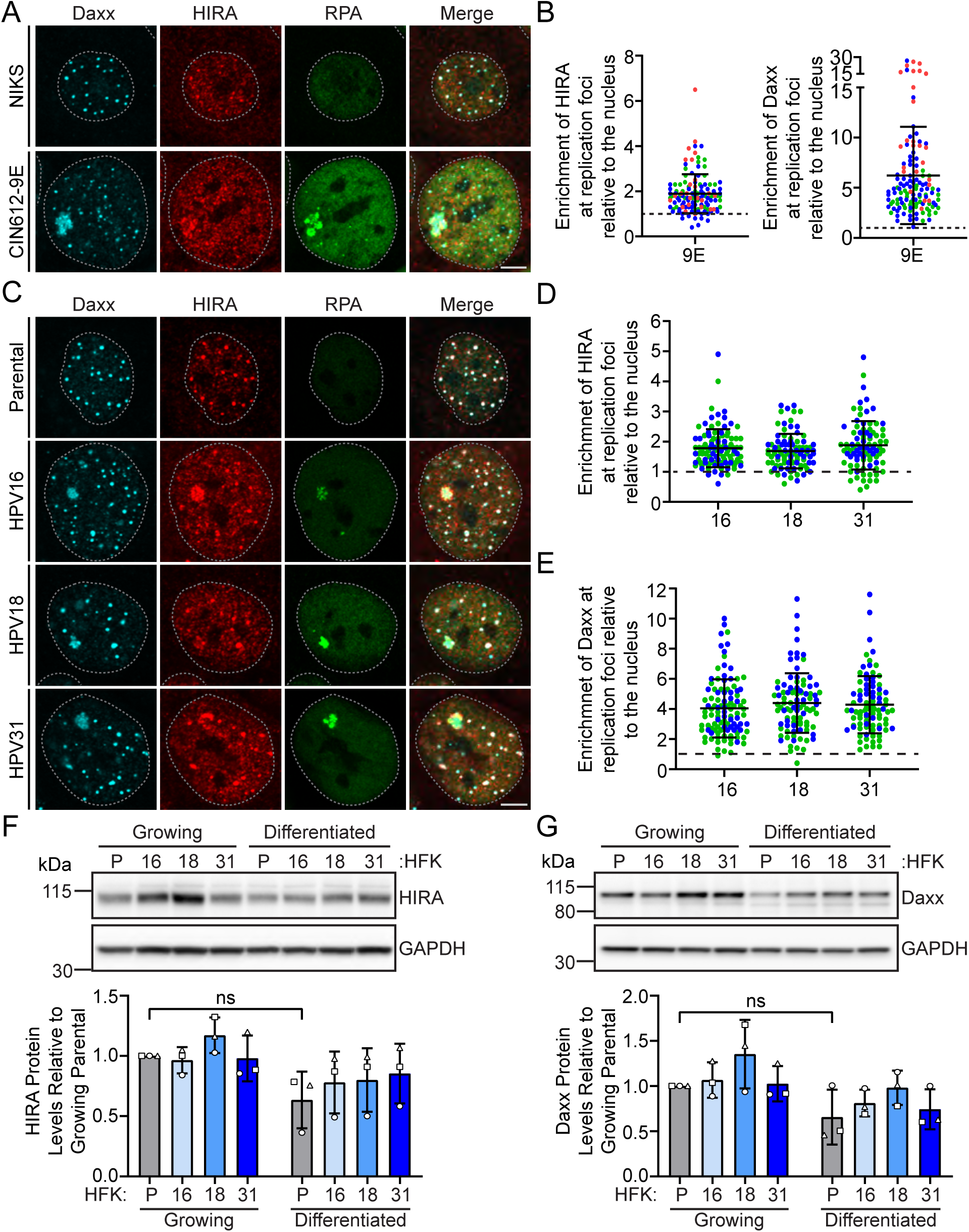
Histone H3.3 chaperones HIRA and Daxx localize to replication foci in keratinocytes containing various HR-HPV genomes. **A**. Representative immunofluorescence staining of Daxx (cyan), HIRA (red) and RPA (green) in differentiated NIKS and 9E cells (n=3). **B**. The raw integrated density of HIRA and Daxx at the HPV replication foci was calculated using ImageJ (values > 1.0 denote enrichment over surrounding nucleoplasm) (n=3; minimum of 25 replication foci/20 nuclei per experiment). **C**. Representative immunofluorescence staining of Daxx (cyan), HIRA (red), and RPA (green) in differentiated HPV negative HFK, and HFK cells containing HPV16, HPV18, and HPV31 genomes (n=2). The raw integrated density of HIRA (**Panel D**) and Daxx (**Panel E**) at the HPV replication foci in differentiated HFK cells was calculated using ImageJ (values > 1.0 denote enrichment) (n=2; minimum of 33 replication foci/30 nuclei per experiment). Representative immunoblots of growing and differentiated HPV negative HFK, and HFK cells containing HPV16, HPV18, and HPV31 genomes, visualizing protein levels of HIRA (**Panel F**), Daxx (**Panel G**), and GAPDH (n=3). Quantitation below, showing the protein levels of HIRA or Daxx normalized to GAPDH relative to growing uninfected HFK parental cells (n=3). Shapes represent independent experiments. (n=3). For all images, a gray dotted line outlines the nucleus defined by DAPI staining. Scale bar, 5 μm. Circles denote individual replication foci; colored circles represent independent experiments. Error bars represent ± standard deviation of the mean.

To assess whether HIRA or Daxx protein levels changed with keratinocyte differentiation or viral infection, protein lysates prepared from the matched HFK/HPV cell lines were analyzed by immunoblot. This showed that both HIRA and Daxx levels were unaffected by viral infection or cellular differentiation (**Figure 1F and G**). Therefore, the association of HIRA and Daxx with HPV replication factories is due to relocalization of each protein and not to increased protein expression. We continued to characterize the function of HIRA, but not Daxx, during the late phase of the HPV life cycle because downregulation of Daxx by siRNA impeded keratinocyte differentiation (**Supplementary Figure 1**).

### 2.2 Keratinocyte differentiation induces localization of HIRA to PML-NBs

HIRA is normally distributed throughout the nucleus, but under conditions of stress (such as viral infection, senescence, or interferon treatment) it is observed at PML-NBs [16]. In differentiated HPV-negative NIKS and parental HFKs HIRA localized with Daxx, a constituent protein of PML-NBs, in punctate nuclear bodies that resembled PML-NBs (**Figure 1A and C**). Immunofluorescence staining for PML, HIRA, and RPA confirmed that these nuclear bodies were PML-NBs (**Supplementary Figure 2A and C**). HIRA also associated with PML-NBs both throughout the nucleus and at sites of HPV replication in HPV genome containing cells (**Supplementary Figure 2**). HIRA localization to PML-NBs greatly increased following calcium induced differentiation of uninfected NIKS keratinocytes (**Supplementary Figure 3**). Therefore, keratinocyte differentiation is sufficient to induce localization of HIRA to PML-NBs, in addition to the previously described promotion by viral infection, interferon treatment or cellular senescence [16, 20].

### 2.3 Spatial organization of HIRA in HPV31 replication foci

Many cellular factors are recruited to HPV replication foci [21–26] and they are often spatially distinct within these factories [19]. To determine whether HIRA localized to specific regions or processes within the HPV31 replication foci, differentiated 9E cells were stained with antibodies specific to RNA Polymerase II (active RNA Pol II phosphorylated on serine residue 2), RPA, and Rad51, or labeled with EdU (5-ethynyl-2’deoxyuridine) to identify regions of transcription, DNA replication, DNA recombination, or nascent DNA synthesis, respectively. Cells were analyzed by confocal imaging and 3D image reconstruction.

As shown in **Figure 2A-B**, active transcription (defined by RNA Pol II) localized to the surface of HPV31 replication foci with little or no colocalization with HIRA (**Supplementary Figure 4A**). To ascertain whether HIRA localized to sites of nascent HPV31 DNA synthesis within the replication foci, EdU was incorporated into newly synthesized DNA for one hour prior to fixation of differentiated 9E cells. As shown in **Figure 2C-D**, HIRA localized to actively replicating HPV31 replication foci but was not directly colocalized with all EdU signals (**Supplementary Figure 4B**). Therefore, HIRA was partially but not exclusively localized to nascently replicating DNA.

**Figure 2:**
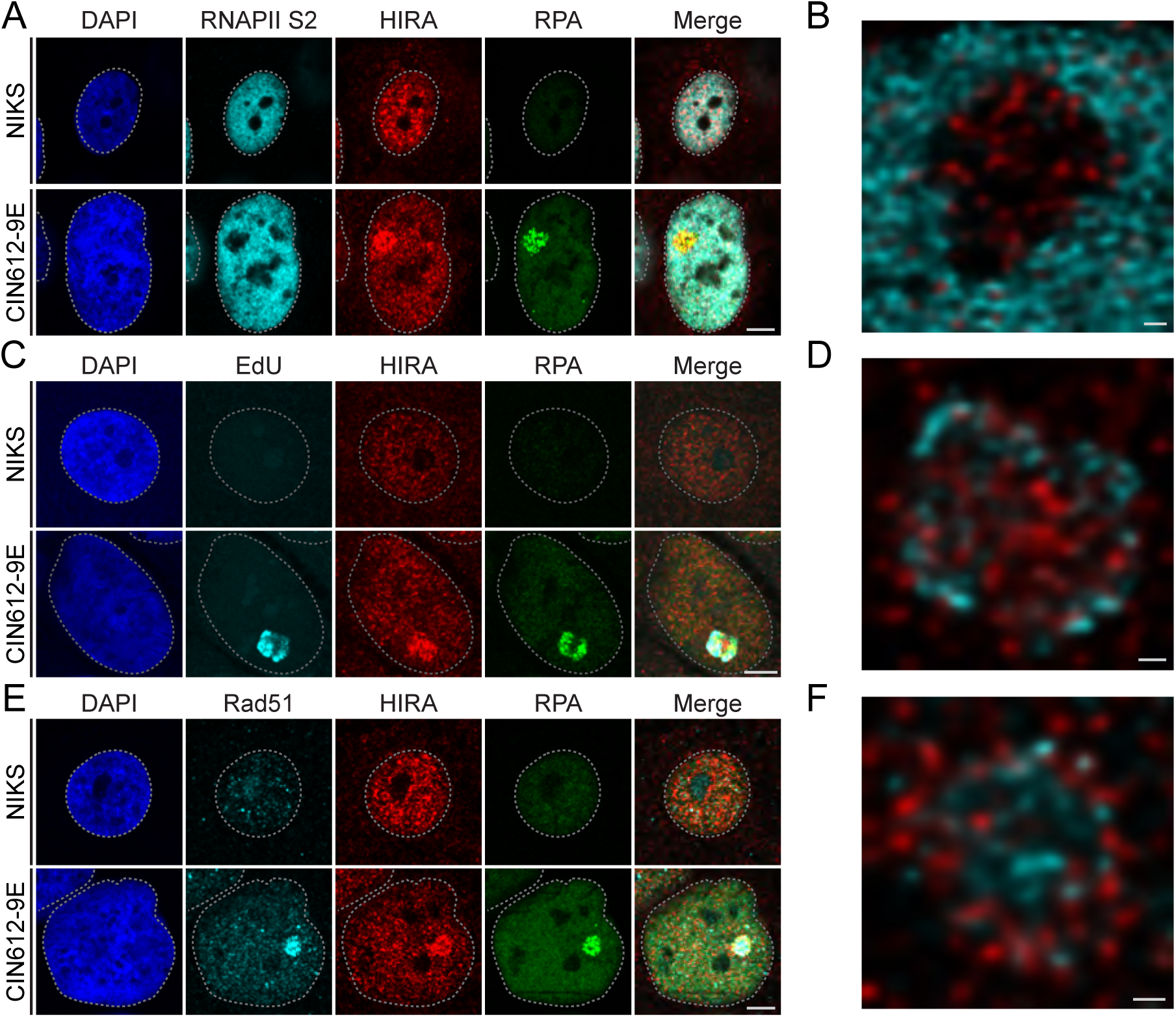
HIRA spatial organization in keratinocytes containing HPV31 genomes. **A**. Representative immunofluorescence staining of RNAPIIS2 (cyan), HIRA (red), and RPA (green) in differentiated NIKS and 9E cells from three independent experiments. **B**. A high-resolution image from a single optical slice of a deconvolved Z-stack, showing RNAPIIS2 (cyan) and HIRA (red) within the replication foci from Panel A. **C**. Representative immunofluorescence staining of EdU (pseudo-colored cyan), HIRA (pseudo-colored red), and RPA (pseudo-colored green) in differentiated NIKS and 9E cells from two independent experiments. **D**. A high-resolution image from a single optical slice of a deconvolved Z-stack, showing EdU (cyan) and HIRA (red) within the replication foci from Panel C. **E**. Representative immunofluorescence staining of Rad51 (cyan), HIRA (red), and RPA (green) in differentiated NIKS and 9E cells from three independent experiments. **F**. A high-resolution image from a single optical slice of a deconvolved Z-stack, showing Rad51 (cyan) and HIRA (red) within the replication foci from Panel E. For all images, DAPI staining of the nucleus is shown in blue. A gray dotted line outlines the nucleus. Scale bar in Panel A, C, and E, 5 μm. Scale bar in Panel B, D, and F, 0.5 μm.

HPV DNA replication also requires factors from the DNA damage response (DDR) pathway [21, 25, 27, 28] and productive replication of viral genomes involves recombination-directed replication processes [23, 29]. As such, we determined whether HIRA localization correlated with the single stranded DNA binding protein RPA, or the Rad51 recombinase involved in homologous recombination [26]. As shown in **Figure 2 and Supplementary Figure 4**, HIRA localized throughout the replication foci, in a pattern that was similar to, but also distinct from, that of RPA. HIRA and Rad51 were also enriched at most replication foci (**Figure 2E-F**) with only a moderate overlap (**Supplementary Figure 4C**).

Altogether, these data indicate that HIRA localization at HPV replication foci does not correlate exclusively with any one function such as transcription or replication; rather HIRA may function simply as a chaperone to load histones onto viral DNA during several processes.

### 2.4 HIRA associates with replication foci formed transiently by E1 and E2 protein expression

Analysis of HPV replication foci in differentiated keratinocytes provides the ideal environment to study wild type HPV DNA replication, but it is not ideal for genetic analysis because many viral mutations are deleterious to the complete infectious cycle. Alternatively, HPV replication foci can be generated in undifferentiated keratinocytes by transient expression of the E1 and E2 replication proteins in the presence of an HPV genome or a plasmid containing the minimal replication origin. Large replication foci form within 24 hours and recruit cellular factors similar to the foci that form in differentiated keratinocytes [24]. Therefore, to determine whether HIRA is recruited to E1-E2 foci, HPV16 E1 and E2 expression plasmids were transfected into HFK cells along with pUC19 negative control plasmid DNA (-ORI), a plasmid containing the HPV16 origin of replication (+ORI), or the full HPV16 genome (+Genome). As shown in **Figure 3**, this resulted in the formation of nuclear foci as detected by the EE-tagged HPV16 E1 protein. These foci show strong enrichment of both HIRA and RPA in foci containing the origin plasmid (>75% foci per nuclei showed HIRA recruitment). We conclude that replication foci generated by amplification of the minimal origin plasmid by the E1 and E2 viral proteins can recruit HIRA and that no other viral proteins are required. When cells were transfected with an HPV genome and the E1 and E2 expression plasmids, the foci formed were much larger and concomitantly were strongly enriched with HIRA in ∼80% foci per nucleus. This suggests that HIRA is directly involved in viral DNA replication. In the absence of any viral replicon, smaller E1-E2 foci formed, and these had only partial and weak association with HIRA in about 40% of foci per nuclei (**Figure 3**). As shown previously, these foci represent DNA damage resulting from E1 expression [28]. Collectively, these data support a role for HIRA in viral DNA replication.

**Figure 3:**
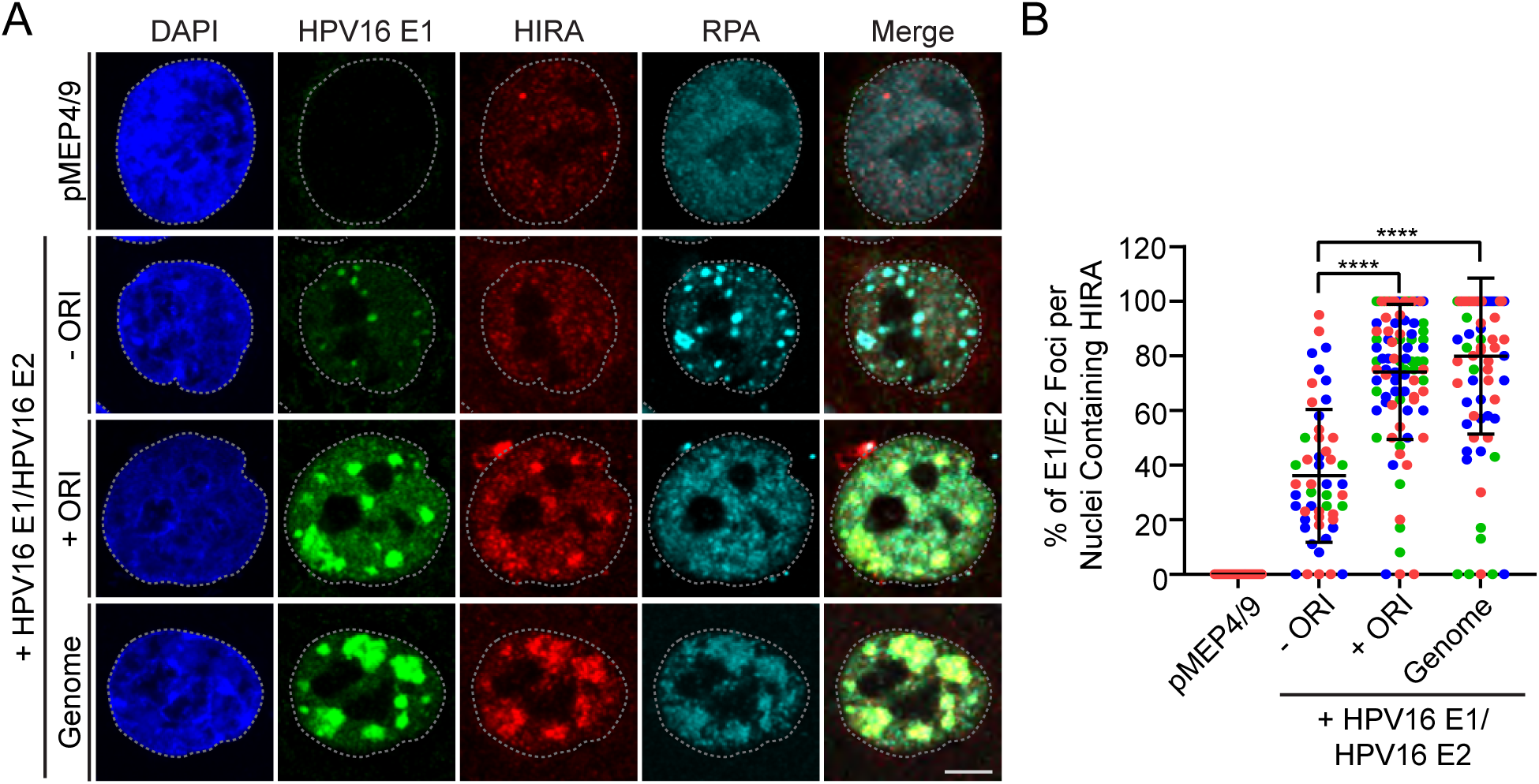
HIRA localization to HPV16 E1/E2 nuclear foci does not require other HPV proteins other than replication proteins E1 and E2. **A.** Representative immunofluorescence staining of EE-tagged HPV16 E1 (green), HIRA (red), and RPA (cyan) in conditionally immortalized HFK cells transfected with HPV16 E1/E2 expression plasmids from three independent experiments. DAPI staining of the nucleus is shown in blue. A gray dotted line outlines the nucleus. Scale bar, 5 μm. **B.** The percentage of replication foci per nuclei where HIRA is enriched at HPV16 E1/E2 replication factories marked by HPV16 E1 (n=3); scored a minimum of 6 nuclei [pMEP4/9], 7 nuclei/81 replication foci [E1/E2/-ORI], and 30 nuclei/160 replication foci [E1/E2/+ORI or Genome]). Circles denote individual replication foci; colored circles represent independent experiments. Error bars represent ± standard deviation of the mean and statistical significance was calculated using an unpaired Mann-Whitney test. ****, P ≤ 0.0001.

### 2.5 HIRA association with HPV16 replication foci occurs in a E8^E2-independent manner

Replication and transcription of HPV genomes are tightly regulated by the HPV E8^E2 repressor protein [30], and keratinocytes containing HPV16 E8^E2 deficient genomes have increased viral DNA copy number and increased early and late viral transcription [31]. Our laboratory has shown that keratinocytes containing HPV16 genomes defective in E8^E2 expression (E8-Mutant) form numerous, large HPV replication foci in undifferentiated cells and recruit many factors similar to wildtype HPV16 genomes [19]. The H2A variant histone macroH2A and the viral E8^E2 repressor protein (in complex with NCoR/SMRT cellular transcriptional repressors) and are both recruited to wildtype HPV replication foci [19, 32]. HPV replication foci are greatly increased in size in the absence of the E8^E2 repressor protein and macroH2A and NCoR/SMRT proteins are absent. If HIRA is important for delivering histones to replication foci for viral DNA replication, we would expect to see an enrichment of HIRA in these foci. In contrast, if HIRA is directly associated with the E8^E2 protein, macroH2A, or NCoR/SMRT proteins, we would expect to observe depletion of HIRA. As shown in **Figure 4**, HIRA is enriched in the large replication foci in keratinocytes containing HPV16 E8 mutant genomes. Therefore, we conclude that neither the viral E8^E2 protein, nor the host macroH2A or NCoR/SMRT proteins are required for this association.

**Figure 4:**
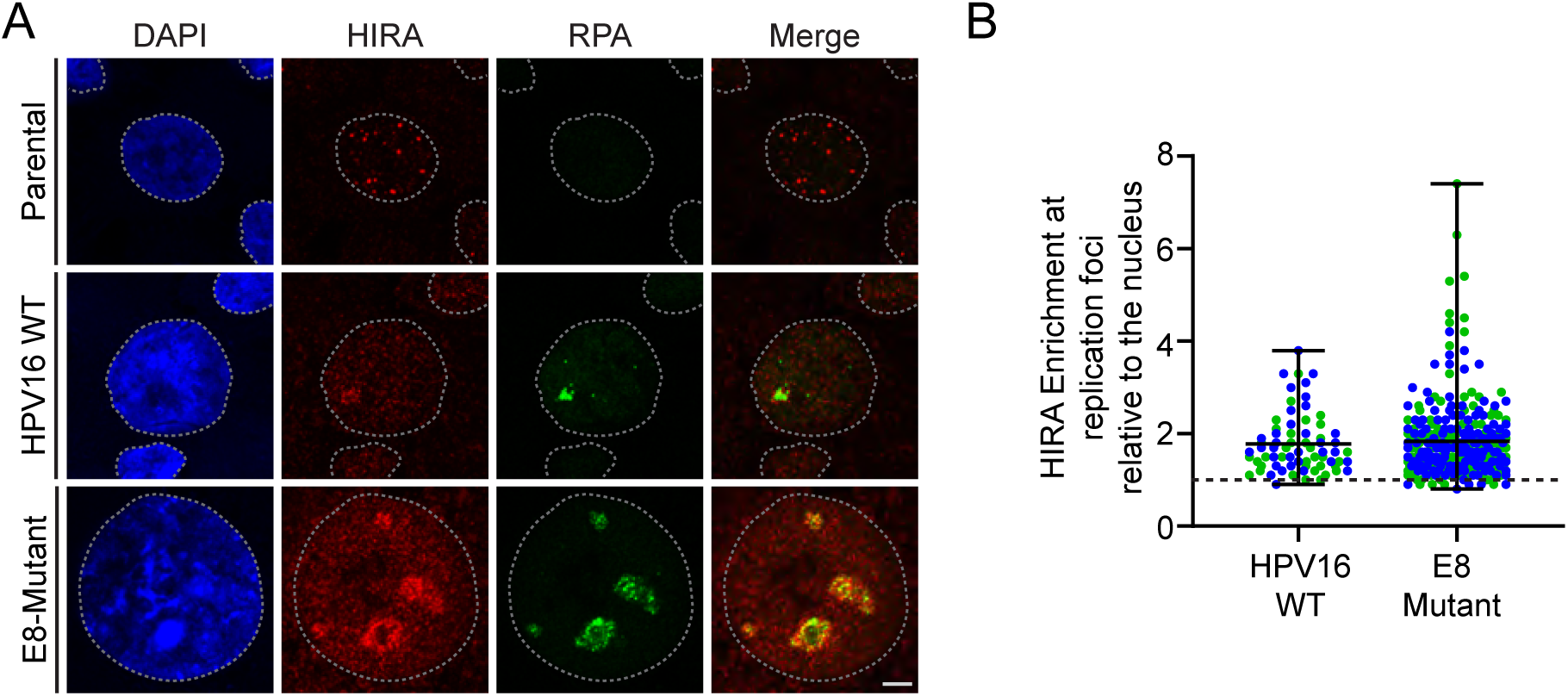
HIRA associates with HPV16 replication foci in an E8^E2-independent manner in primary keratinocytes. **A**. Representative immunofluorescence staining of HIRA (pseudo-colored red) and RPA (green) in differentiated HFK39 cells with and without HPV16 genomes and growing HFK39 cells containing HPV16 E8-Mutant genomes from two independent experiments. DAPI staining of the nucleus is shown in blue. A gray dotted line outlines the nucleus. Scale bar, 5 μm. **B**. The raw integrated density of HIRA at replication foci was calculated using ImageJ (values > 1.0 denote enrichment) [n=2]. Circles denote individual replication foci, colored circles represent independent experiments; measured a minimum of 34 replication foci/30 nuclei per experiment. Error bars represent the range surrounding the mean.

### 2.6 HIRA localization to sites of HPV31 DNA replication does not require Sp100

Our laboratory has previously shown that Sp100 is observed adjacent to and infiltrated in HPV replication factories in cells containing extrachromosomal HPV genomes where it restricts replication and transcription [33]. We and others have also shown that HIRA localizes to PML-NBs in a Sp100-dependent manner [16, 34, 35], but the interplay of Sp100 and HIRA at HPV replication foci is unknown. Given that Sp100 is required for HIRA localization to PML-NBs and is present at sites of HPV replication [36] we evaluated whether Sp100 was required for HIRA localization to HPV replication foci.

To assess the requirement of Sp100 in the localization of HIRA to sites of HPV replication, we downregulated Sp100 levels in differentiated 9E cells using siRNA transfection and assessed HIRA localization by immunofluorescence. Cells were transfected with pools of control (siCtrl) or Sp100 (siSp100) as outlined in the scheme in **Figure 5A**. Sp100 depletion was moderate (∼2.2 fold) at the protein level and did not perturb HIRA protein levels (**Figure 5B-C**). However, depletion of Sp100 resulted in decreased mRNA levels of all four Sp100 isoforms (**Figure 5D**). As shown in **Figure 5E**, Sp100 was observed adjacent to and sometimes speckled throughout HPV31 replication foci, while HIRA was enriched throughout the majority of HPV31 replication foci observed in siCtrl cells (**Figure 5F**). Foci containing cells treated with Sp100 siRNA did not contain detectable Sp100, but this did not alter HIRA localization to the HPV31 replication foci (**Figure 5E-F**). These results, taken together, show that Sp100 does not regulate HIRA protein levels or influence HIRA localization to HPV replication foci.

**Figure 5:**
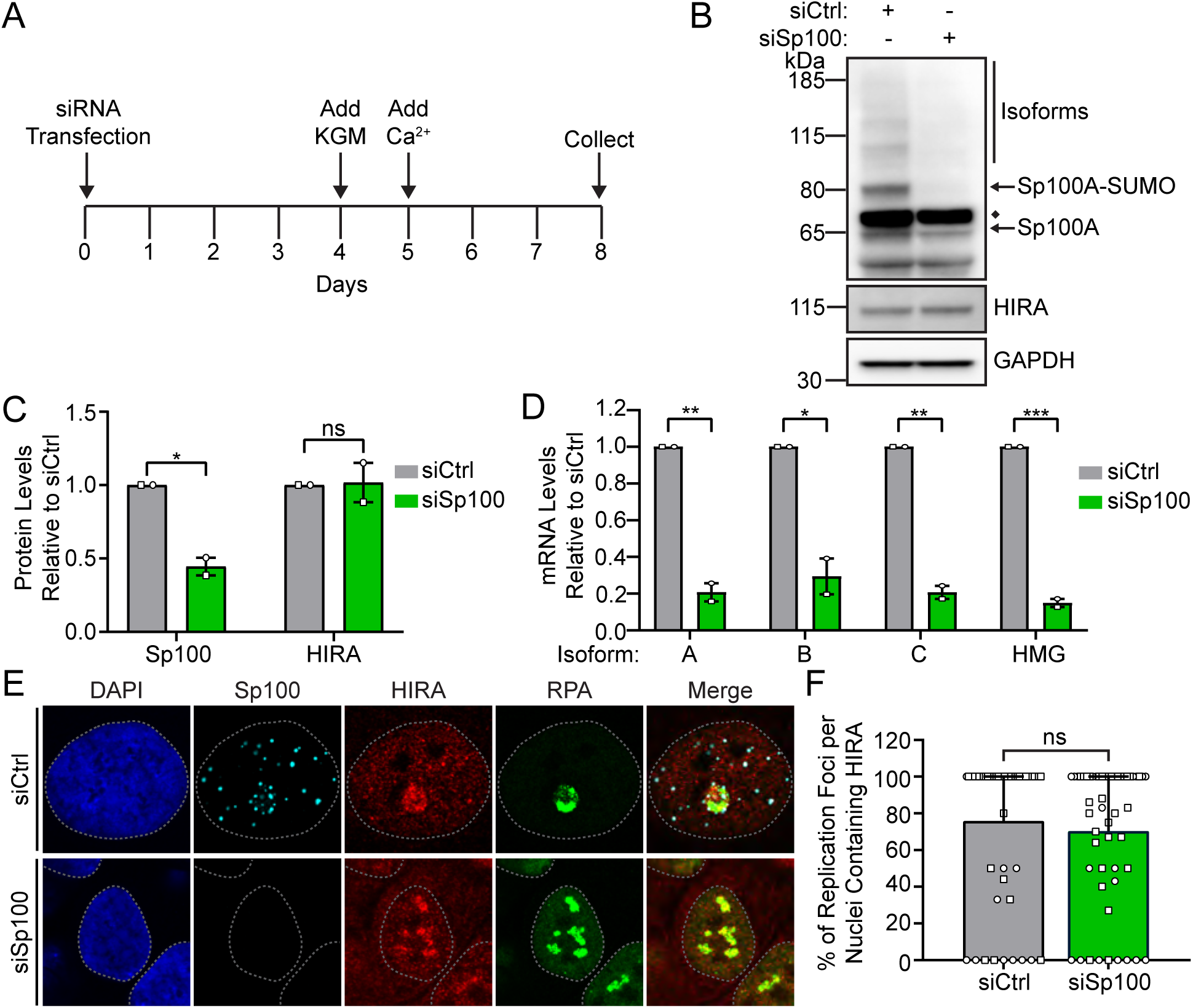
HIRA localization to HPV31 replication foci occurs in a Sp100-independent manner. **A**. Timeline of siRNA knockdown in 9E cells. Cells were plated at low density and transfected with 25 nM control (siCtrl) or Sp100 (siSp100) siRNA after 24 hours. Differentiated cells were collected 8 days post transfection. **B.** Representative immunoblots of differentiated 9E siCtrl and siSp100 cells, visualizing the protein levels of Sp100, HIRA, and GAPDH (n=2). The single GAPDH blot shown is representative of both blots but only the one matched to the Sp100 blot is shown. **C.** Quantitation of panel B, showing the fold change in total Sp100 and HIRA protein levels normalized to GAPDH relative to siCtrl (n=2). **D.** qPCR measuring Sp100 transcripts, showing the fold change in Sp100 mRNA levels normalized to PPIA relative to siCtrl (n=2). **E.** Representative immunofluorescence staining of Sp100 (cyan), HIRA (red), and RPA (green) in differentiated 9E cells transfected with siCtrl or siSp100 from two independent experiments. DAPI staining of the nucleus is shown in blue. A gray dotted line outlines the nucleus. Scale bars, 5 μm. **F.** The percentage of replication foci per nuclei where HIRA is localized to HPV31 replication factories marked by RPA (n=2; scored a minimum of 22 nuclei/32 replication foci per condition). Shapes represent independent experiments. Shown are the mean values and error bars represent the range. Statistical significance was calculated using an unpaired student’s t-test (C-D) or Mann-Whitney test (F). ns, not statistically significant, *, P ≤ 0.05, **, P ≤ 0.01, *** P ≤ 0. 001.

### 2.7 HPV31 DNA amplification is modestly reduced by depletion of HIRA

To directly assess the requirement of HIRA for viral DNA replication, we downregulated HIRA levels in 9E cells using siRNA and measured viral DNA levels in undifferentiated and differentiated cells by Southern blot analysis and qPCR. Cells were transfected with control (siCtrl), or HIRA (siHIRA) siRNA pools and total cellular DNA was isolated at day two (growing) or day eight (differentiated) post-transfection, as shown in the scheme in **Supplementary Figure 5A**. Depletion of HIRA was efficient both at the protein (≥ ∼ 2.9 fold decrease) and mRNA (≥ ∼ 5.7 fold decrease) level **(Supplementary Figure 5B-D)**. The calcium differentiation method does not consistently result in amplification of viral DNA and so we only evaluated experiments that did. The number of HPV31 DNA copies per cell were modestly reduced upon HIRA depletion (∼2.5 fold) in differentiated cells as measured by Southern blot and qPCR (**Figure 6A-C**). Analysis of replication foci as detected by RPA immunofluorescence in differentiated 9E cells showed that downregulation of HIRA did not perturb replication foci formation (**Supplemental Figure 5E-F**). These findings show that HIRA modestly promotes levels of late HPV31 DNA amplification.

**Figure 6:**
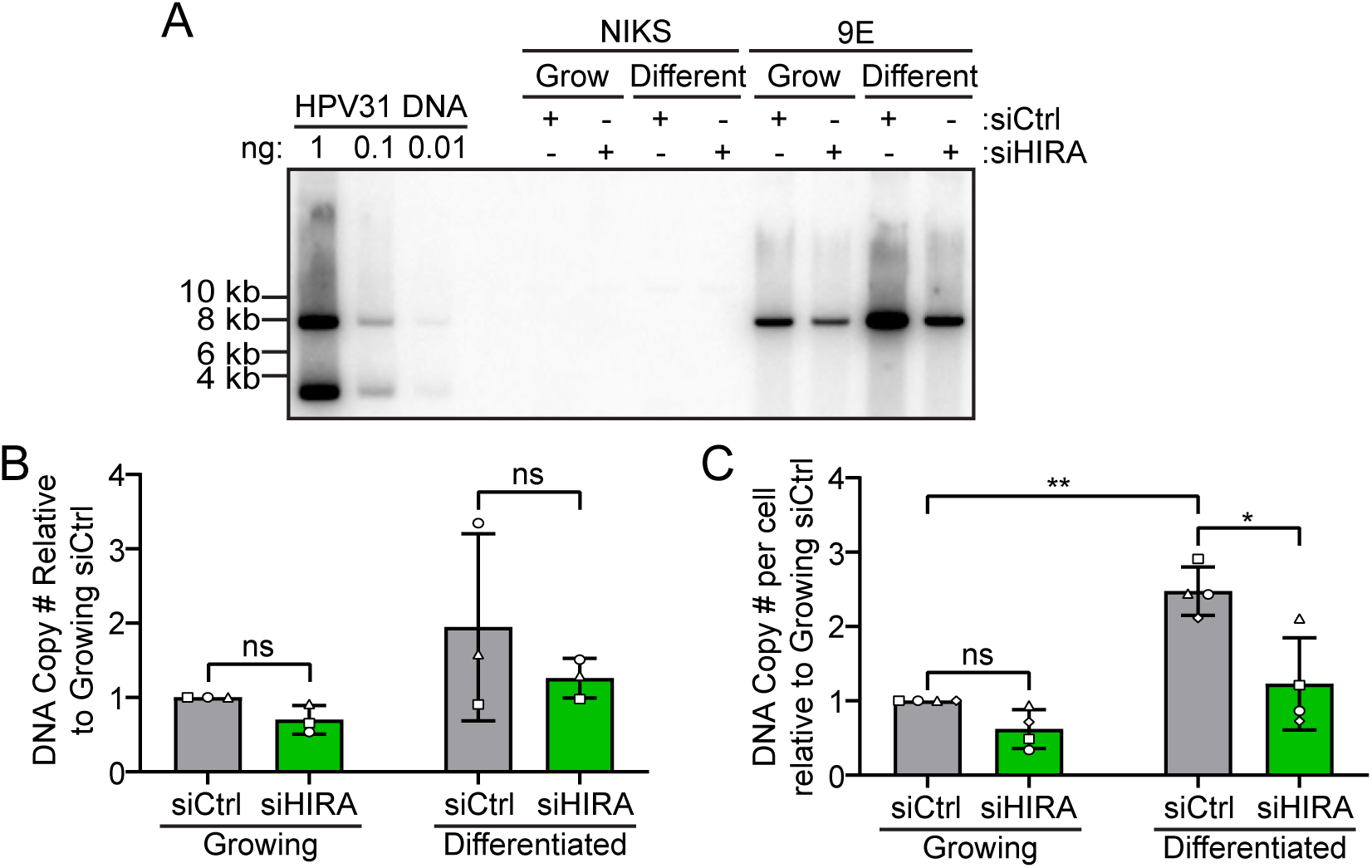
HPV31 DNA amplification is modestly reduced in the absence of HIRA. **A**. Southern blot analysis of DNA extracted from growing and differentiated NIKS and 9E cells transfected with siCtrl or siHIRA, showing linearized HPV31 DNA. DNA copy number controls for 1, 0.1, and 0.01 ng of HPV31 DNA are shown. **B.** Quantitation of panel A, showing the fold change in linear HPV31 DNA in 9E cells relative to siCtrl (n = 3). **C.** DNA qPCR for HPV31 DNA using DNA extracted from growing and differentiated 9E cells transfected with siCtrl or siHIRA, showing the fold change in HPV31 DNA copies per cell relative to growing siCtrl in cells containing amplified HPV31 genomes (n = 4). Shapes represent independent experiments. Error bars represent ± standard deviation of the mean and statistical significance was calculated using a paired student’s t-test. ns, not statistically significant, *, P ≤ 0.05, **, P ≤ 0.01.

### 2.8 HIRA modestly enhances late HPV31 transcription

To determine whether HIRA is important for viral transcription, we downregulated the levels of HIRA in 9E cells and measured different spliced viral transcripts shown in **Figure 7A** in undifferentiated and differentiated cells by qRT-PCR. Cells were transfected with control (siCtrl) or HIRA (siHIRA) siRNA and total cellular RNA was isolated at two (growing) or eight days (differentiated) post-transfection, similar to the scheme in **Supplementary Figure 5A**. We found no difference in E6* (early) or E1^E4 (intermediate) transcript levels in undifferentiated cells (**Figure 7**). As expected, both E1^E4 and L1 transcripts were greatly increased by differentiation. Downregulation of HIRA did not alter early or intermediate viral transcripts **(Figure 7B-C**) in differentiated cells but resulted in a modest (though statistically insignificant) reduction in the late L1 viral transcript (**Figure 7D**).

**Figure 7:**
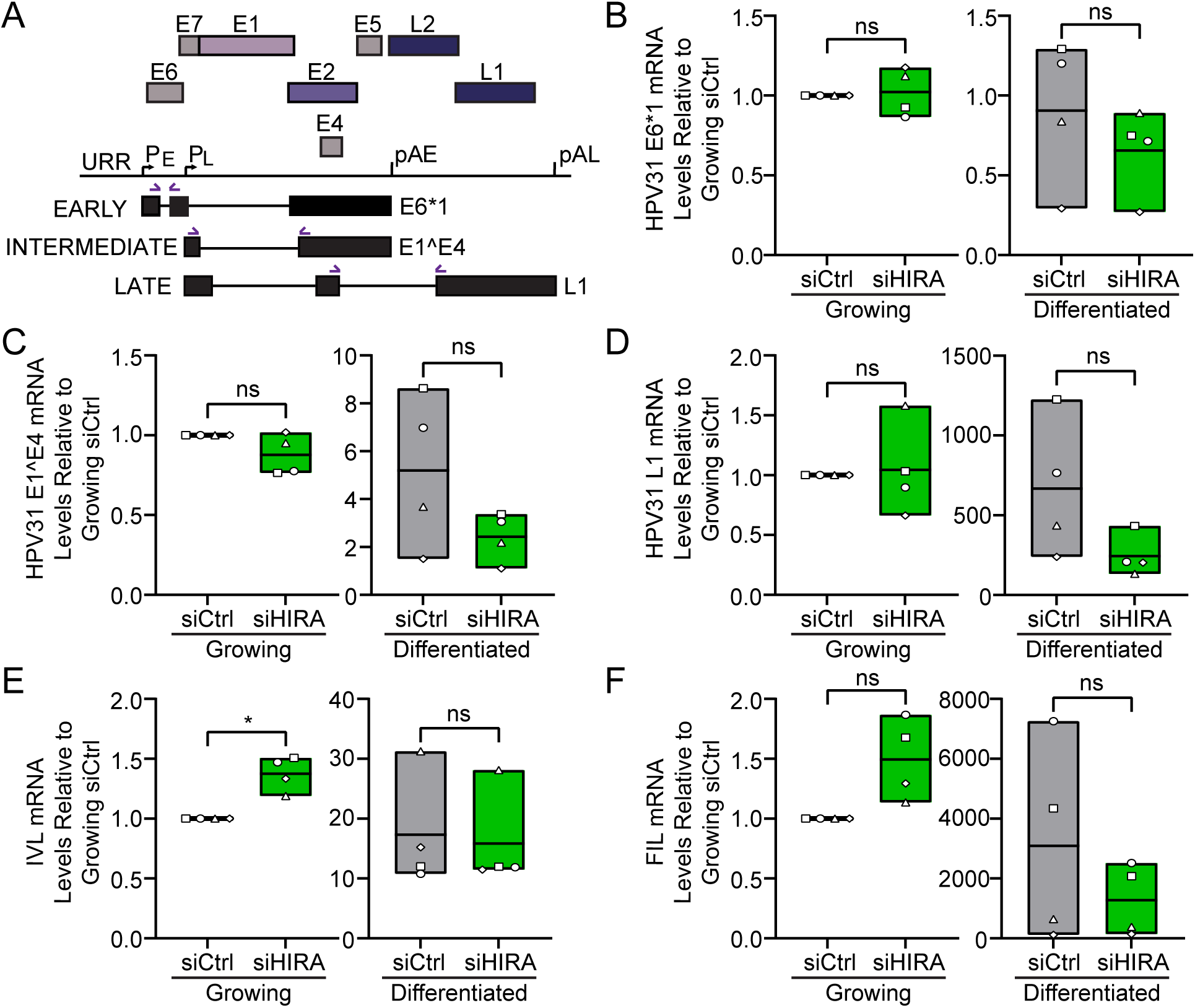
Late HPV31 transcription is modestly reduced following depletion of HIRA. **A**. Diagram of the HPV31 viral transcripts and qRT-PCR strategy used to detect viral spliced mRNAs. Spliced primer pairs are denoted with purple arrows. **B-D.** qRT-PCR using reverse-transcribed RNA from growing and differentiated 9E cells transfected with siCtrl or siHIRA, showing the fold change in HPV31 E6*1, E1^E4, and L1 viral transcripts normalized to PPIA within its own condition relative to growing siCtrl (n = 4). **E-F.** qRT-PCR using reverse-transcribed RNA from growing and differentiated 9E cells transfected with siCtrl or siHIRA, showing the fold change in involucrin and filaggrin cellular transcripts normalized to PPIA within its own condition relative to growing siCtrl (n=4). RNA was isolated 2 days and 8 days post transfection for the growing and differentiated conditions, respectively. Floating bars represent minimum and maximum values, and the line indicates the mean. Shapes represent independent experiments. Statistical significance was calculated using a paired student’s t-test. ns, not statistically significant, *, P ≤ 0.05.

Late viral transcription is dependent on the host epithelial differentiation program; therefore, we measured the mRNA levels of two cellular differentiation markers involucrin and filaggrin, to determine if downregulation of HIRA influenced host keratinocyte differentiation. As shown in **Figure 7E and F**, depletion of HIRA did not significantly alter mRNA levels of involucrin but resulted in a modest reduction in filaggrin in differentiated cells. We conclude that while there is a modest decrease in late transcription in the absence of HIRA supporting a role for HIRA in the promotion of late HPV31 viral transcription, this could also be due to the decrease in viral genome copy numbers detected in **Figure 6A-C** or to reduced keratinocyte differentiation.

### 2.9 HIRA Complex factors are enriched in HPV replication foci

HIRA functions as a histone H3.3 chaperone in complex with calcineurin-binding protein 1 (CABIN1), Ubinuclein 1 (UBN1) and transiently interacts with the Anti-silencing function 1A (ASF1a) histone chaperone [3]. HIRA, UBN1 and ASF1a, directly associate with H3.3 to facilitate deposition of histone H3.3 mediated by HIRA [37]. Therefore, we analyzed HPV31 replication foci for the presence of these factors by immunofluorescence. Differentiated HPV negative NIKS cells were used as an uninfected control. As shown in **Figure 8A-D**, UBN1 and ASF1a, and HIRA, were associated with PML-NBs in both uninfected and HPV31 containing cells, and in most HIRA positive replication foci in differentiated 9E cells. Taken together, the presence of the HIRA complex members in replication foci indicates that HIRA is likely to be functioning as a histone chaperone for productive viral DNA replication.

**Figure 8:**
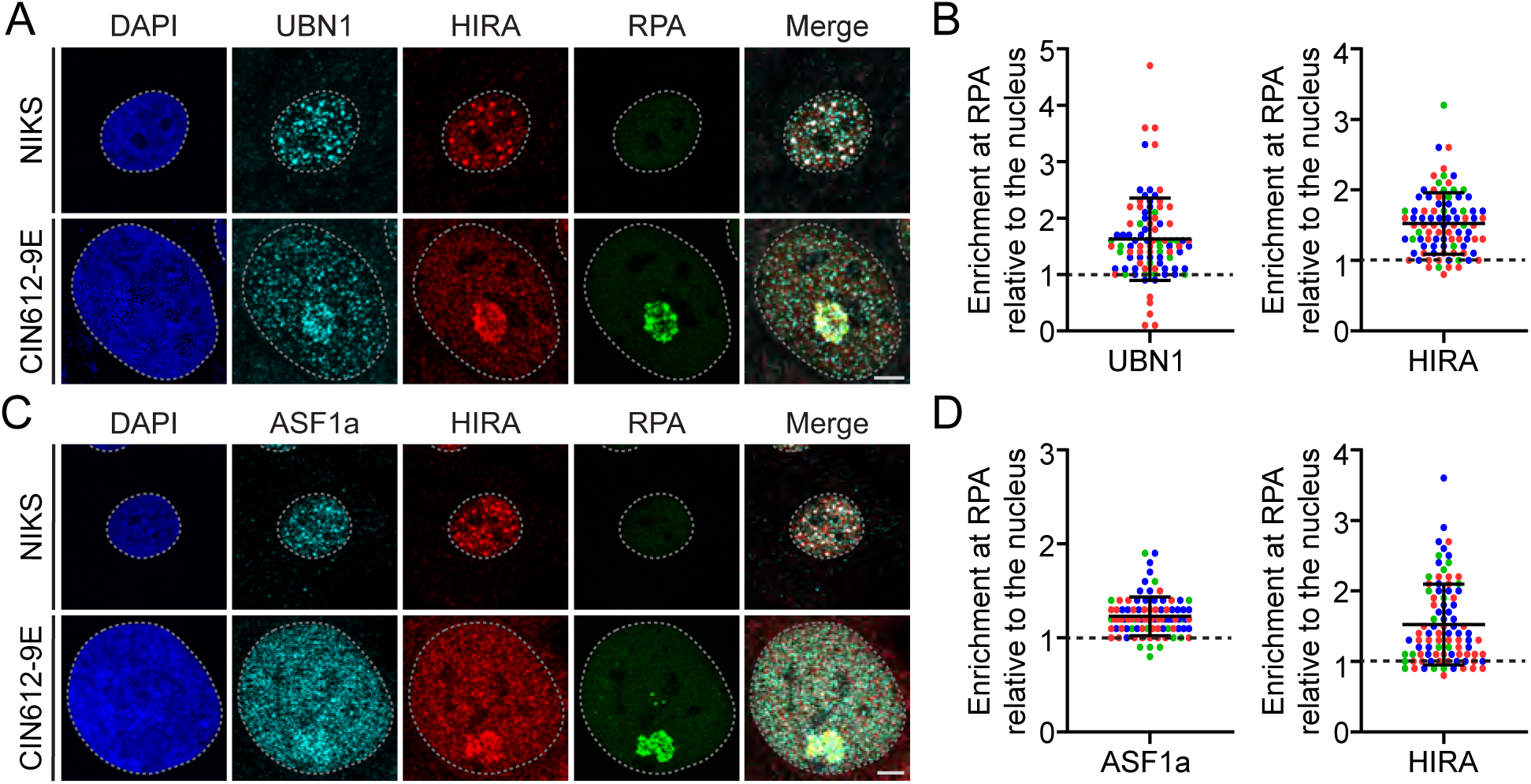
The histone H3.3 chaperone HIRA and associated complex members, UBN1 and ASF1a, localize to late replication foci in keratinocytes containing HPV31 genomes. Representative immunofluorescence staining of UBN1 [**Panel A**] or ASF1a [**Panel C**] (pseudo-colored cyan), HIRA (pseudo-colored red), and RPA (green) in differentiated NIKS and 9E cells from three independent experiments. For all images, a gray dotted line outlines the nucleus defined by DAPI staining. Scale bar, 5 μm. The raw integrated density of UBN1 and HIRA [**Panel B**] and ASF1a and HIRA [**Panel D**] at the HPV replication foci in differentiated 9E cells was calculated using ImageJ (values > 1.0 denote enrichment) n = 3; minimum of 16 replication foci/10 nuclei per experiment. Circles denote individual replication foci; colors represent independent experiments. Error bars represent ± standard deviation of the mean.

### 2.10 Histone H3.3 is the predominant H3 histone in differentiated keratinocytes

HIRA is a histone H3.3 chaperone and its presence at sites of HPV replication implies deposition of histone H3.3 onto viral genomes. The canonical histone H3.1 is only expressed in S-phase [1], and so H3.3 is predicted to be the major H3 histone in differentiated keratinocytes. Furthermore, HPV DNA amplification occurs in a G2-like phase of the cell cycle in these cells [5, 6]. H3.1 is encoded by ten genes derived from the HIST1 gene cluster and the H3.3 variant is encoded by two separate genes, H3F3A and H3F3B (**Supplementary Figure 6A**). We recently conducted an RNAseq analysis of total RNA from proliferating and differentiated cells from the HFK/HPV cell lines shown in **Figure 1C and Supplementary Figure 2C** (the complete analysis will be published elsewhere). As shown in **Supplementary Figure 6B**, there was a great decrease in H3.1 transcripts after differentiation of parental HFK cells, while H3.3 transcripts remained at constant levels. Therefore, H3.3 is most abundant during replication of HPV minichromosomes in differentiated keratinocytes.

### 2.11 Histone H3.3 phosphorylated on serine 31 is associated with HPV replication foci

To determine if histone H3.3 is enriched at sites of viral replication, we examined the association of histone H3.3 with viral replication foci formed in differentiated 9E cells by immunofluorescence. All commercially available H3.3 antibodies tested did not detect a nuclear pattern consistent with histones (they gave rise to a speckled nuclear pattern) and as shown in **Figure 9A-B**, H3.3 was not enriched at HPV31 replication foci using these antibodies.

**Figure 9:**
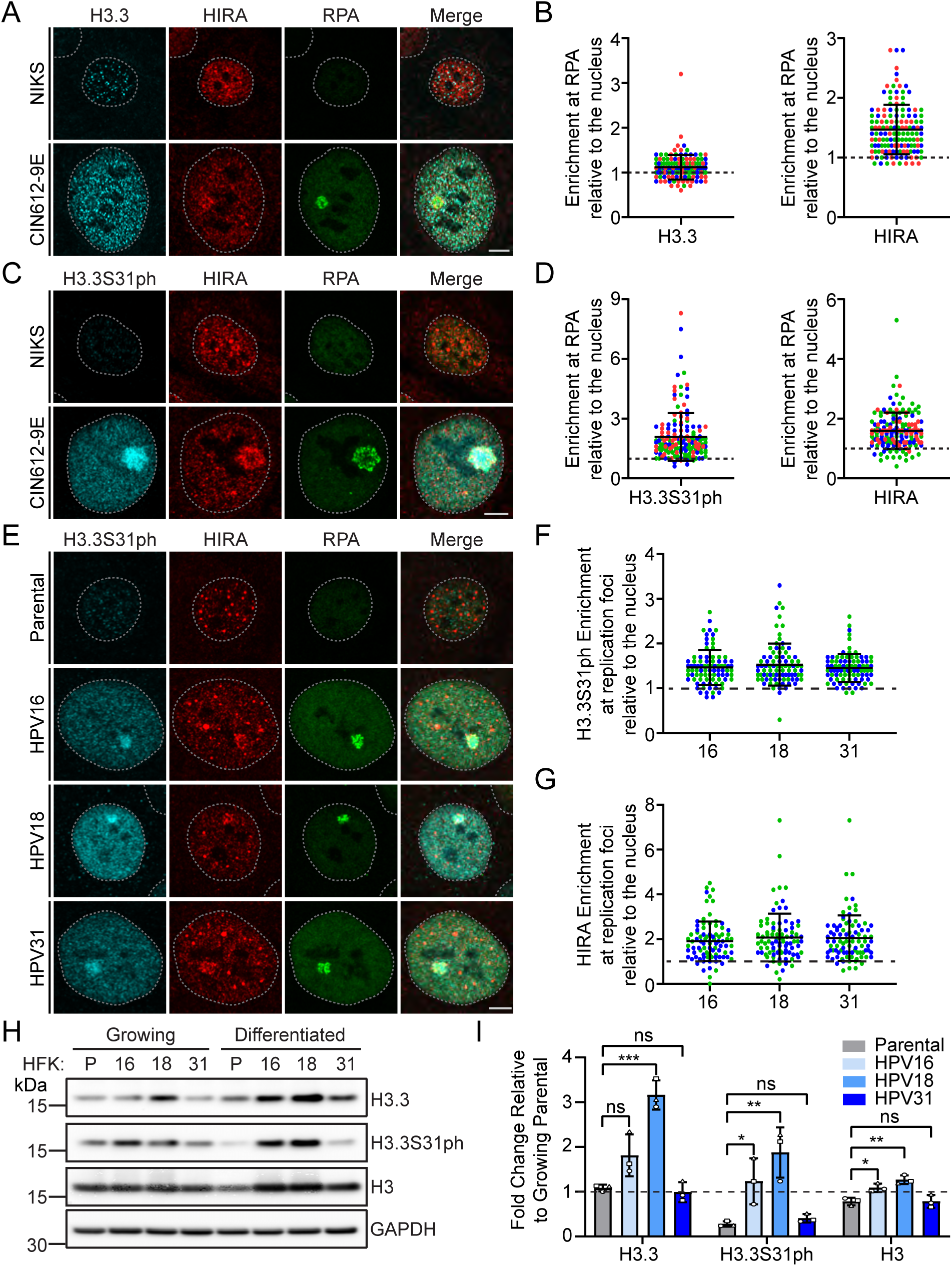
Histone variant H3.3 phosphorylated at serine 31 is enriched at HPV replication foci in keratinocytes containing various HR-HPV genomes. **A**. Representative immunofluorescence staining of H3.3 (pseudo-colored cyan), HIRA (pseudo-colored red), and RPA (green) in differentiated NIKS and 9E cells from three independent experiments. **B**. The raw integrated density of H3.3 and HIRA at the foci in differentiated keratinocyte cells was calculated using ImageJ (values > 1.0 denote enrichment). **C**. Representative immunofluorescence staining of H3.3S31ph (cyan), HIRA (red), and RPA (green) in differentiated NIKS and 9E cells from three independent experiments. **D**. The raw integrated density of H3.3S31ph and HIRA at the foci in differentiated keratinocyte cells was calculated using ImageJ (values > 1.0 denote enrichment). **E**. Representative immunofluorescence staining of H3.3S31ph (cyan), HIRA (red), and RPA (green) in differentiated HPV negative HFK, and HFK cells containing HPV16, HPV18, and HPV31 from two independent experiments. The raw integrated density of H3.3S31ph [**Panel F**] or HIRA [**Panel G**] at the foci in differentiated keratinocyte cells was calculated using ImageJ (values > 1.0 denote enrichment). For all images, a gray dotted line outlines the nucleus defined by DAPI staining. Scale bar, 5 μm. **H**. Representative immunoblots of growing and differentiated HPV negative HFK, and HFK cells containing HPV16, HPV18, and HPV31 genomes, visualizing the protein levels of H3.3, H3.3S31ph, H3, and GAPDH (n=3). The GAPDH blot shown is representative of all blots, but matched specifically to that of the H3.3 blot. **I**. Quantitation of panel H, showing the protein levels of H3.3, H3.3S31ph, and H3 in differentiated cells normalized to GAPDH relative to growing uninfected HFK parental [black dotted line] (n=3). Shapes represent independent experiments. For quantitation panels B and D (n=3) and panels F and G (n=2); Circles denote individual replication foci; colored circles represent independent experiments. Scored a minimum of 39 replication foci/30 nuclei per experiment. Error bars represent ± standard deviation of the mean and statistical significance was calculated using an unpaired student’s t-test. ns, not statistically significant, *, P ≤ 0.05, **, P ≤ 0.01, ***, P ≤ 0.001.

To explore this further, we analyzed the location of H3.3 histone phosphorylated on serine residue 31 (H3.3S31ph), a post-translational modification unique to H3.3 and associated with both stimulation-induced transcription [38] and promotion of heterochromatin formation [39]. Using an antibody specific for H3.3S31ph, we observed that, in contrast to H3.3, H3.3S31ph was greatly enriched in the majority (89%) of HPV31 replication factories in 9E cells **(Figure 9C-D**). H3.3S31ph localization also occurred at most replication foci formed by HPV16, HPV18 and HPV31 (**Figure 9E-G**). Immunoblot analysis showed that histone H3, H3.3 and H3.3S31ph were present at variable levels in growing and differentiated cells containing the different HPV genome, but no consistent effect was observed due to differentiation or the presence of HPV (**Figure 9H-I**). These data indicate that histone H3.3 deposited onto HPV DNA during late amplification of the viral genome, is highly phosphorylated on serine 31.

### 2.12 Enrichment of histone variant H3.3 phosphorylated at serine 31 at HPV31 replication foci is independent of HIRA

To determine whether H3.3S31ph association was dependent on HIRA, we downregulated HIRA expression with siRNA and examined replication foci in 9E cells for the presence of H3.3S31ph. As shown in **Supplementary Figure 7**, HIRA and H3.3S31ph were enriched at the majority (96%) of HPV31 replication foci in siCtrl treated cells, however downregulation of HIRA did not perturb the enrichment (96%) or intensity of H3.3S31ph at HPV replication foci. To determine whether downregulation of HIRA influenced protein levels of H3.3 and H3.3S31ph, protein levels were analyzed by western blot. The total H3.3 protein levels, but not the modified H3.3S31ph, were slightly decreased following depletion of HIRA (**Supplementary Figure 7D-E**). We conclude that HIRA is not required for H3.3 serine 31 phosphorylation in HPV replication foci. Unfortunately, we could not determine whether deposition of H3.3 itself was reduced in replication foci after downregulation of HIRA because of the aforementioned issue with H3.3 antibodies.

### 2.13 Proposed role of H3.3 deposition during HPV replication and its link to late HPV transcription

H3.3S31ph is enriched at sites of HPV replication (**Figure 9**) and this modification could be important to the processes that take place in viral replication factories. Of note, the Chk1 kinase is central to the process of HPV DNA replication and can phosphorylate H3.3 on residue 31 [40]. HPV DNA amplification requires activation of both the ataxia telangiectasia mutated (ATM) and ATM and Rad3-related (ATR) arms of the DNA damage response [27, 41, 42] and these pathways can be induced by E7 and/or E1 expression [25, 27, 28]. E7 induces phosphorylation of STAT5, which increases transcription of the scaffold protein, TopBP1 [43]. TopBP1 localizes to sites of HPV replication where it is required for the initiation of HPV replication [44]; TopBP1 also directly binds to RPA coated single-stranded DNA at sites of DNA damage, and in turn, activates the ATR/Chk1 pathway [45]. Activation of the Chk1 kinase is essential for HPV DNA amplification and its inhibition abrogates HPV DNA amplification [43]. Phosphorylation of H3.3 serine residue 31 results in the recruitment and activation of the histone acetyl transferase p300, which acetylates histone H3 on residue 27 (H3K27ac) resulting in increased Brd4 binding [38, 46–48]. HPV replication foci have been shown to contain acetylated histones and Brd4, which modulates both viral DNA replication and transcription [19, 24, 49–54]. These pathways are outlined in **Figure 10A**.

**Figure 10:**
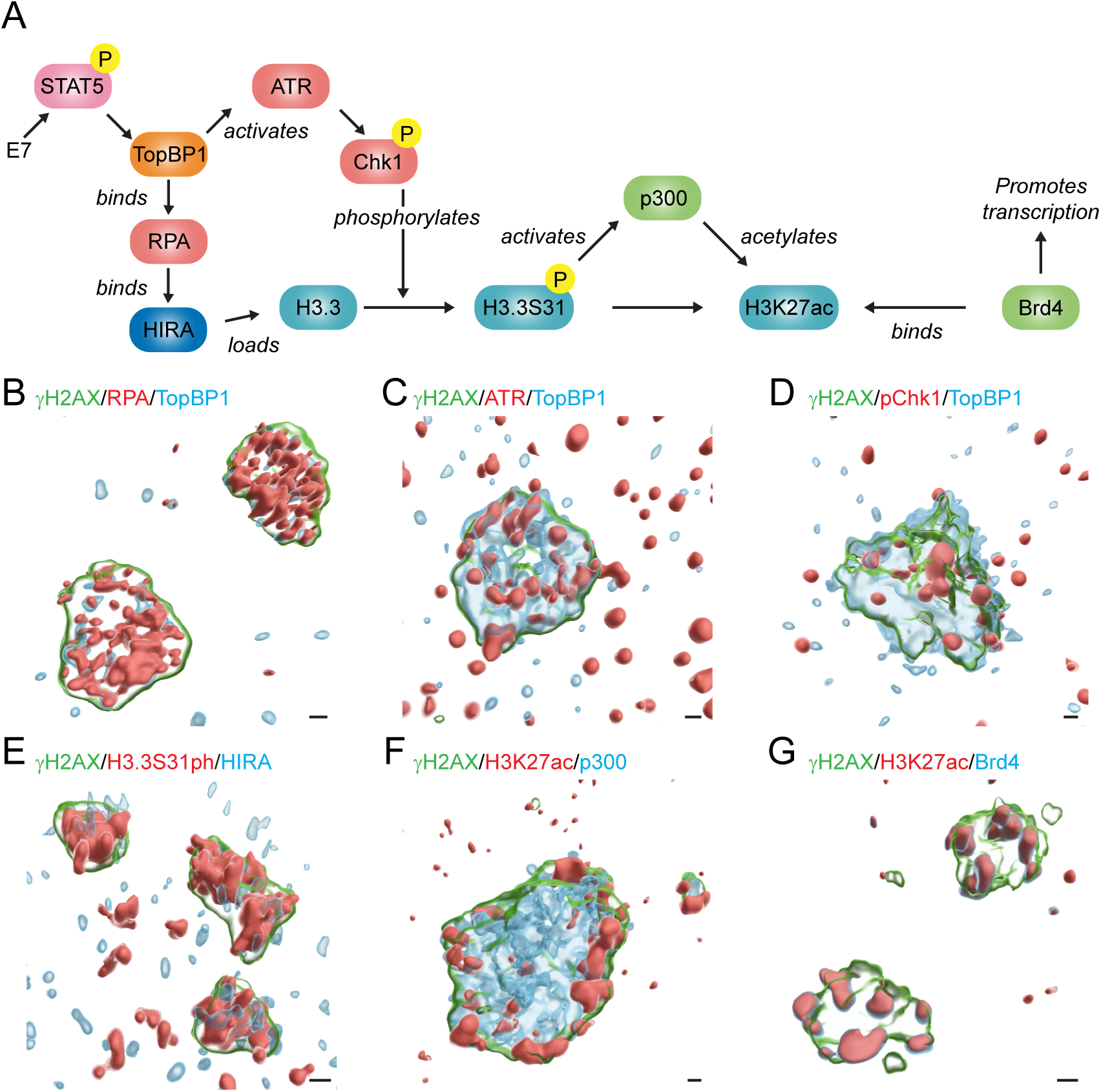
Proposed role of H3.3 deposition during HPV replication and its link to late HPV transcription. **A.** Schematic of the proposed pathway in which HPV replication occurs through activation of the host DDR through the ATR/Chk1 pathway, leading to the phosphorylation of the histone variant H3.3, and how this phosphorylation impacts the process of transcription. Z-Stack images were processed in Huygens Essential (cross-talk correction and deconvolution) and surface renderings were created in Imaris to visualize HPV31 replication foci stained for **B.** γH2AX (green), RPA (red), and TopBP1 (cyan), **C.** γH2AX (green), ATR (red), and TopBP1 (cyan), **D.** γH2AX (green), pChk1 (red), and TopBP1 (cyan), **E.** γH2AX (green), H3.3S31ph (red), and HIRA (cyan), **F.** γH2AX (green), H3K27ac (red), and p300 (cyan), and **G.** γH2AX (green), H3K27ac (red), and Brd4 (cyan) in differentiated 9E cells. Scale bars, 0.5 μm. All images shown are derived from cells fixed in 4% PFA in PBS, but nuclei shown in Panel B-C and Panels E-G were incubated in CSK buffer prior to fixation.

Figure 10 shows representative 3D graphical images of these factors in HPV31 replication factories. Differentiated 9E cells were stained with specific antibodies and HPV31 replication foci were examined by confocal imaging and 3D image reconstruction. In each case, the replication foci were visualized using an antibody against γH2AX to delineate the chromatin of replication foci. As shown in Figure 10B-C, TopBP1, RPA, and ATR were enriched inside the foci at sites of active HPV31 replication and/or recombination. ATR and Chk1 are activated in HPV containing cells but these signaling molecules are transient and not consistently tightly bound to DNA damaged viral or host chromatin (**Figure 10E**) [56, 57]. RPA can regulate the deposition of H3.3 onto DNA through direct binding to HIRA [55]. HIRA and H3.3S31ph were observed localized throughout the replication foci (**Figure 10E**). As shown previously [19, 24], p300, H3K27ac and Brd4 were found on the surface of the replication foci (Figure 10F-G) and are thought to be important for viral transcription. Altogether, we propose that HPV activation of the ATR pathway leads to the incorporation of H3.3, its subsequent phosphorylation on serine 31, and downstream activation of viral transcription.

## 3. Discussion

### 3.1 Overview

In this study we characterize the role of HIRA, a histone H3.3 chaperone, that is enriched at replication factories during productive HPV DNA replication. HIRA functions independently from host DNA replication and instead deposits nucleosomes at active, histone-depleted, regions of chromatin. Productive HPV replication is unconventional; large amounts of DNA are synthesized in a short period of time in an unlicensed recombination-directed replication (RDR) process that relies heavily on the DNA damage response and homologous replication. Viral genome replication must be coordinated with high levels of late viral transcription to generate the L1 and L2 capsid proteins. This process occurs in differentiated cells that express mostly H3.3 compared to H3.1 and so the histone chaperone HIRA is likely required to support multiple viral chromatin-associated processes.

### 3.2 HIRA localizes to PML-NBs in differentiated keratinocytes

This project was originally inspired by a study by McFarlane *et al*. that showed that infection by an ICP0 deficient mutant of HSV1 induced localization of HIRA to PML-NBs as part of an anti-viral innate immune response [16]. We examined this phenomenon in keratinocytes and discovered that HIRA was observed at PML-NBs in both uninfected and HPV containing keratinocytes. HIRA has previously been shown to localize to PML-NBs during viral infection, cellular senescence or after interferon treatment and here we show that this localization can also be promoted by keratinocyte differentiation. We surmise that this could be due to increased innate immune signaling resulting from the calcium differentiation protocol [58] or due to senescence-associated pathways in the primary keratinocytes [20]. The remainder of our study focused on the observed localization of HIRA at replication factories.

### 3.3 The role of the PML-NB component Sp100 in HIRA localization

We have recently shown that the Sp100 component of PML-NBs is required for HIRA localization to these bodies in keratinocytes [35]. We have also shown that Sp100 localizes to sites of HPV31 replication to restrict late viral transcription and replication [33]. We show here that HIRA localizes to sites of HPV replication independently from Sp100 and promotes late viral replication and transcription. Thus, we conclude that these host factors have opposing roles at late stages of the HPV life cycle.

### 3.4 Spatial analysis of HIRA and cellular factors in HPV replication foci

Numerous cellular factors are recruited to, and have different spatial locations within, HPV replication foci to facilitate viral genome replication and transcription. To determine whether HIRA was associated with one specific process, we used high-resolution microscopy to investigate the relative location of HIRA and other factors in HPV replication foci. We have shown previously that RNA polymerase localizes to the surface of the foci [19], and Ray-Gallet and colleagues have demonstrated an interaction between HIRA and RNA Pol II at transcription sites [59]. However, spatial analysis of HIRA and RNA Pol II showed minimal overlap within the replication foci, suggesting that HIRA localization at HPV replication foci is not solely to promote viral transcription. Instead, we observed HIRA throughout the foci in moderate association with nascently synthesized DNA and host DDR factors known to facilitate HPV DNA replication. These results indicate that HIRA does not function exclusively in any singular process occurring at sites of HPV replication. The HIRA associated complex members UBN1 and ASF1a also localize to sites of HPV31 replication. As HIRA, UBN1, and ASF1a are capable of binding with H3.3-H4 dimers [37], this strongly implies that deposition of H3.3 is occurring at sites of HPV replication.

### 3.5 The role of viral and host proteins in recruitment of HIRA to replication factories

We show that HIRA is highly enriched in foci formed by amplification of a minimal viral origin containing replicon by expression of the E1 and E2 proteins. We conclude that no other viral proteins are required for this recruitment. HIRA partially localized to a small percentage of E1-E2 foci that form in the absence of viral DNA due to DNA damage of host DNA [28]. This localization is not unexpected as HIRA mediated deposition of H3.3 occurs at damaged chromatin during DNA repair and subsequent recovery of transcription [8]. The HPV E8^E2 repressor protein restricts high levels of viral genome amplification and gene expression in undifferentiated cells [30]. Previous work from our laboratory indicated that both the NCoR/SMRT transcriptional repressor proteins and histone variant macroH2A are excluded from HPV replication foci generated by a viral genome defective in E8^E2 expression [19]. Here we found that HIRA was highly enriched in replication foci formed by the E8^E2 mutant genome.

These results indicate that HIRA functions independently of the NCoR/SMRT complex, macroH2A, and E8^E2. In addition, the strong enrichment of HIRA took place in undifferentiated cells that should contain histone H3.1 and associated chaperones, further supporting the specificity for HIRA as the preferred histone chaperone. In the experiments described herein, HIRA enrichment strongly correlated with increased viral DNA replication and focus formation.

### 3.6 Phosphorylated H3.3 is highly enriched in HPV replication foci

Replication and transcription during the late phase of the viral lifecycle relies on the exploitation of the host DDR [7] and multiple components of the host ATR/Chk1 DDR pathway are localized at sites of HPV31 replication. We show that the phosphorylated version of H3.3, H3.3S31ph, is enriched at replication factories of multiple HPV types. This H3.3 specific post-translational modification links the ATR DNA damage response to highly active chromatin [46, 47], resulting in histone acetylation and recruitment of Brd4 that could promote late viral transcription and other viral chromatin-associated processes. Chk1 is one of three kinases that phosphorylate serine 31 [46, 60] and is absolutely required for HPV DNA amplification in differentiated cells [43].

Porter and colleagues have shown that HPV minichromosomes packaged in virions are enriched in histone H3.3 and it was proposed that this could be a consequence of replication within differentiated cells. It was also hypothesized that this could also be a strategy to prime the genome for early transcription and replication upon infection of the next cell [4]. Future studies could assess whether H3.3S31ph is incorporated into viral chromatin and impacts infection of a new host cell.

### 3.7 Technical Limitations of our study

Our study has several technical limitations. RNAseq analysis confirmed that histone H3.3, rather than H3.1, was the most abundant histone H3 in differentiated cells, but we were unable to convincingly detect H3.3 by immunofluorescence using commercial antibodies. Future studies are required to establish a direct interaction of H3.3 with the viral genome and to discern if this association is dependent on HIRA. Meanwhile, we used an antibody against phosphoserine 31, a residue specific to the H3.3 histone variant, which proved to be a useful and interesting substitute. Another caveat is that depletion of HIRA did not reduce H3.3S31ph enrichment at HPV31 replication foci. It has been reported that depletion of H3.3 is only accomplished by reduction of both H3.3 chaperones, HIRA and Daxx [11]. Daxx also associates with HPV replication foci and promotes HPV18 and HPV11 DNA replication in U2OS cells [61]. Lynskey and colleagues show that HIRA mediated deposition of H3.3 prevents the accumulation of single-stranded DNA devoid of nucleosomes in the absence of Daxx [60]. However, in our system, depletion of Daxx levels greatly decreased keratinocyte differentiation thus preventing downregulation of both HIRA and Daxx in differentiated keratinocytes. It is plausible that Daxx also deposits H3.3 at HPV replication foci in the absence of HIRA, and this putative potential compensatory mechanism would underscore the importance of H3.3 deposition during late HPV replication.

To determine the role of HIRA during the infectious cycle, we used siRNA transfection to reduce HIRA levels. Depletion of HIRA resulted in a two-fold decrease in HPV31 DNA replication and L1 transcription, suggesting that HIRA is a positive regulator during the late phase of the HPV life cycle. However, downregulation of HIRA also modestly reduced expression of the keratinocyte differentiation marker, filaggrin. Therefore, decreased L1 transcription could also result indirectly from decreased HPV DNA replication or from reduced keratinocyte differentiation.

Depletion of HIRA did not perturb formation of HPV31 replication foci and viral DNA synthesis and histone deposition are likely uncoupled; The large amounts of single-stranded DNA present at sites of HPV replication (as detected by RPA staining) could reflect this uncoupling and could also explain the affinity for HIRA at HPV replication factories. It is probable that replication continues with depletion of HIRA, but nucleosome occupancy on replicating HPV genomes is reduced.

The Chk1 kinase is central to the pathway that we propose in Figure 10. However, chemical inhibition of the ATR/Chk1 pathway abrogates HPV DNA amplification in differentiated cells [43] and so long term-experiments were not feasible.

### 4.0 Conclusions

Our findings that an H3.3 chaperone localizes to and promotes HPV replication highlights a novel way in which HPV takes advantage of the cellular environment and processes to enable viral replication in differentiated keratinocytes. We propose that the key function of HIRA is to chaperone and load histone H3.3 onto replicating viral DNA. Concomitant engagement of the ATR arm of the DNA damage response results in H3.3 phosphorylation on serine 31 by Chk1. In turn, this histone modification promotes transcription by activation of the histone acetylase p300, acetylation of H3K27, and recruitment of Brd4. Therefore, we reveal a novel way in which HPVs hijack multiple processes to promote the late phases of replication in differentiated keratinocytes.

## 5. Materials and Methods

### 5.1 Plasmids

Metallothionein inducible Glu-Glu (EE) tagged HPV16 E1 (pMEP9:EEHPV16 E1 Recoded) and Flag tagged HPV16 E2 (pMEP4:FlagHPV16E2 Recoded2X), and pMEP4, and pMEP9 expression vectors were described previously [28]. The HPV16 minimal origin of replication plasmid (pKS-HPV16ori) was previously described [62], in addition to the HPV16 minicircle genome [63] The DNA standard curves for qPCR were generated from the 9E HPV31 genome in HindIII of pBR322 [64] or the pMA-T HPV31 E6*1, pMA-T HPV31 E1^E4, pCR2.1 HPV31 L1 spliced 3590^5552, pMT-Sp100 isoform, pCR2.1-TopoTA involucrin, pCR2.1-TopoTA filaggrin, and pCR2.1-TopoTA cyclophilin A plasmids [33]. The Ribonuclease P RNA component H1 (RPPH1) DNA standard curve was generated from the pMA RPPH1 gene plasmid (Thermo Fisher; GeneArt) and the HIRA DNA standard curve was generated from the pCMV-SPORT6-HIRA plasmid (Horizon Discovery; MGC Human HIRA Sequence-verified cDNA).

### 5.2 Cell Lines

A cell line derived from a low-grade cervical intraepithelial neoplasia containing HPV31 (CIN612-9E) [65] and Nearly diploid immortal keratinocytes (NIKS) cells [66] Have been previously described. Human foreskin keratinocytes (HFKs) and conditionally immortalized HFKs (HFK1a) were previously isolated from neonatal foreskins [67]. Primary HFK strain 39 cells containing replicating wildtype HPV16 or E8^E2 mutant genomes were generated as previously described [68]. Cell lines containing extrachromosomal HPV16 (Genbank, AF125673.1) [69], HPV18 (Genbank, X05015), or HPV31 (Genbank, PP706107) [64] genomes were established by immortalization of primary HFK strain 20 cells. HPV genomes were released from their vectors by restriction enzyme digestion and recircularized by low concentration ligation. Recircularized genomes were electroporated into HFKs along with a plasmid encoding G418 resistance (pCpGneo) [70]. Cells were selected for six days with 300 µg/ml G418 and subsequently passed without selection until colonies formed. Southern blot analysis was performed to ensure extrachromosomal genomes in each cell line, as described previously [70].

### 5.3 Cell Culture

Keratinocytes were cocultured in Rheinwald-Green F medium (3:1 Ham’s F-12/high-glucose Dulbecco’s modified Eagle’s medium [DMEM], 5% fetal bovine serum [FBS], 0.4 mg/ml hydrocortisone, 8.4 ng/ml cholera toxin, 10 ng/ml epidermal growth factor, 24 µg/ml adenine, 6 µg/ml insulin) with lethally irradiated J2-3T3 murine fibroblasts. J2-3T3 cells were grown in DMEM with 10% bovine calf serum) and exposed to 60 Grays gamma irradiation prior to coculture with keratinocytes. For differentiation, keratinocyte cels were grown in F-medium until confluent and switched to low-calcium keratinocyte growth medium (KBM, Lonza CC-3101 containing growth supplements SingleQuots, Lonza CC-4131). After 24 hours in low calcium media, the cells were grown in KBM basal medium containing 1.5 mM calcium chloride for an additional three days.

### 5.4 Transfection of siRNA

9E cells were seeded onto 6-well, 12-well, or 10 cm plates containing irradiated J2-3T3 cells and cultured overnight. Pools of four siRNA duplexes that target HIRA (Dharmacon; L-013610-00-0010), Sp100 (Dharmacon; L-015307-00-0010), or a non-targeting control (Dharmacon; D-001810-10-20) were complexed with lipofectamine RNAiMax transfection reagent (Thermo Fisher) in serum-free media and added to cells at final concentration of 25 nM. The siRNA sequences are detailed in Supplementary Table 1. Cells were incubated with siRNA for two or eight days before harvest.

### 5.5 Transient transfection of E1 and E2 expression plasmids to form replication foci

Conditionally immortalized HFKs (HFK1a) were seeded at a density of 2.86 x 10^4^ cells/cm^2^ onto glass coverslips containing 2.0 x 10^4^ cells/cm^2^ irradiated J2-3T3 cells in F-medium supplemented with 10 µM Rho-kinase inhibitor (Y-27632) and cultured overnight. Cells were transfected with pMEP9-HPV16 E1 and pMEP4-HPV16 E2 expression vectors, or pMEP empty vectors (400 ng each) and 50 ng of pUC19 control plasmid DNA, a plasmid with the HPV16 minimal origin of replication p16ori, or an HPV16 minicircle genome, generated as previously described [19, 24]. Cells were transfected with Fugene 6 according to the manufacturer’s instructions. E1 and E2 protein expression was induced with 3 µM CdSO4 for 4 hours prior to fixation of cells with 4% PFA at 24 hours post transfection.

### 5.6 qPCR for viral DNA copy number

Total cellular DNA was extracted from keratinocytes using the DNeasy Blood and Tissue kit (QIAGEN). 15 ng of DNA was amplified by qPCR with 6 µM primers and detected with SYBR green (Roche) using a QuantStudio 7 Flex Real-Time PCR System (Thermo Fisher). Samples were normalized to Ribonuclease P RNA Component H1 (RPPH1) levels. Copy number was calculated by comparison to a standard curve of HPV31 DNA. The primers are listed in Supplementary Table 2.

### 5.7 Southern blot analysis

Total cellular DNA was extracted from keratinocytes using the DNeasy Blood and Tissue kit (QIAGEN). 1-2 µg total DNA was digested with either an enzyme (HindIII) to linearize the HPV31 genome, or with a non-cutting enzyme (BamHI) to linearize cellular DNA. After digestion, DNA fragments were separated by electrophoresis on 0.8% agarose-TAE gels and subsequently transferred onto Nytran Supercharge membranes using a Turbo Blotter (GE Healthcare). The membranes were UV-crosslinked (120 mJ/cm^2^), dried, and prehybridized in hybridization buffer (3X SSC, 2% SDS, 5X Denhardt’s Solution, and 0.2 mg/ml sonicated salmon sperm DNA) for one hour. The membranes were incubated overnight with 25 ng (^32^P)-dCTP labeled HPV31 DNA probe in hybridization buffer. The ^32^P-radiolabeled probe was generated from a plasmid containing the HPV31 genome using a Random Prime labeling kit (Roche). The membranes were washed in 0.1% SDS/0.1X SSC at 65°C and visualized and quantitated by phosphorimaging on a Typhoon scanner (GE Healthcare).

### 5.8 RNA extraction and qPCR of host and viral transcripts

Prior to isolation, J2-3T3 feeder cells were removed and total RNA was isolated with the RNeasy Mini kit (QIAGEN). RNA concentration was determined using a Nanodrop 1000 spectrophotometer (Life Technologies) or Qubit Fluorometric Quantification (Thermo Fisher). RNA integrity was assessed by capillary electrophoresis on a 2100 bioanalyzer system using RNA 6000 nano kits (Agilent Technologies); all samples had RIN values >8.0. Reverse transcription reactions were carried out with the Transcriptor First-Strand Synthesis kit (Roche). Quantitative real-time qPCR was performed using SYBR Green (Roche) and QuantStudio 7 Flex Real-Time PCR System (Thermo Fisher) in compliance with MIQE guidelines. Cloned cDNA plasmids (with amounts ranging between 2.5 x 10^5^ and 2.5 x 10^-2^ fg) were used to generate standard curves.

Standards for the HPV31 spliced transcripts, E6*1 (nt 186 to 210^413 to 416), E1^E4 (nt 857 to 877^3,292 to 3,296), and L1 (nt 3,562 to 3,590^5552 to 5,554) and cellular genes involucrin, filaggrin, and cyclophilin A were described previously [33]. The HIRA standard curve was generated from the pCMV-SPORT6-HIRA plasmid (Horizon Discovery; MGC Human HIRA Sequence-verified cDNA). HIRA primers span the 294/295 exon-exon junction of HIRA (nt 280 to 299). Primer pairs for host and viral transcripts are listed in Supplementary Table 2. Samples were normalized to Peptidylprolyl Isomerase A (PPIA) levels associated with their respective growth conditions.

### 5.9 Indirect Immunofluorescence

Cells were cultured on acid-washed glass coverslips and fixed with 4% paraformaldehyde (PFA) in 1X phosphate buffered saline (PBS) at room temperature for 15 minutes. Where denoted, coverslips were pre-extracted in cytoskeleton (CSK) buffer (piperazine-*N*,*N*′-bis(2-ethanesulfonic acid) (PIPES) (pH 7), 100 mM NaCl, 300 mM sucrose, 3 mM MgCl2, 1 mM EGTA, 0.5% Triton X-100) for 10 minutes prior to fixation. Fixed cells were permeabilized with either 0.1% or 0.5% Triton X-100 (Sigma) in PBS and blocked in 5% (vol/vol) normal donkey serum (Jackson Immunoresearch). Cells were subsequently incubated with primary antibodies for 1 hour at 37℃ or overnight at 4℃. Primary antibodies used were HIRA [WC119] mouse monoclonal (Millipore, 04-1488; 1:100), RPA32/RPA2 (4E4) rat monoclonal (Cell Signaling, 2208; 1:300), PML (H-238) rabbit polyclonal (Santa Cruz, sc-5621; 1:100), Sp100 rabbit polyclonal (Sigma, HPA016707; 1:500), UBN1 rabbit polyclonal (Sigma, HPA061029; 1:100), ASF1a rabbit monoclonal (Cell Signaling, 2990s; 1:100), RNA Pol II S2 rabbit polyclonal (Abcam, ab-5095; 1:100), Rad51 rabbit monoclonal (Abcam, ab133534; 1:100), H3.3 rabbit monoclonal (Abcam, ab176840; 1:200), H3.3S31ph rabbit monoclonal (Abcam, ab92628-100ul; 1:100), Glu-Glu Epitope Tag rabbit polyclonal (Novus, NB600-354; 1:300), Alexa Fluor 488 H2AX [pS139] mouse (BD Pharmingen, 560445; 1:10), TopBP1 [R1180] rabbit (Boner et. al 2002; 1:500), TopBP1 [B-7] mouse monoclonal (Santa Cruz, sc-271043; 1:100), ATR rabbit polyclonal (Cell Signaling, 2790; 1:100), Phospho-Chk1 (Ser345) [133D3] rabbit monoclonal (Cell Signaling, 2348; 1:200), H3K27ac rabbit polyclonal (Sigma, 07-360; 1:100), p300 [3G230/NM-11] mouse monoclonal (Abcam, ab14984; 1:500), and Brd4 [1F11] mouse monoclonal (from Cheng-Ming Chiang; 1:100).

Also reference **Supplementary Table 3** for antibody information. Cells were washed with 1X PBS and incubated with secondary antibodies for 30 minutes at 37 ℃. Secondary antibodies against the primary antibody species were conjugated with Alexa 488, Alexa 594, Rhodamine Red-X, or Alexa 647 (AffiniPure; Jackson Immunoresearch). Cell nuclei were stained with DAPI, and coverslips mounted to glass slides using Prolong Gold (Life Technologies).

### 5.10 Image Analysis

All images were collected with a Leica TCS-SP8 laser scanning confocal microscope using a x63 oil immersion objective. Images were processed using Leica Application Suite X Software (Leica Microsystems, version 3.7.0.20979), Huygens Essential (Scientific Volume Imaging B.V, version 25.04.0p3 64b), and Imaris (Bitplane, version 10.2.0). Images were exported as tif files and were assembled into figures using Adobe Illustrator (version 28.1).

ImageJ for Enrichment analysis: To measure the enrichment of HIRA and additional target proteins at sites of viral replication, ImageJ (java version;1.8.0_345) was used to define region of interests (ROIs) encompassing the viral replication foci (RPA signal), in addition to regions within the nucleus and cytoplasm. The raw integrated density within the ROIs was measured for the target of interest and then divided by the area of the associated ROI. Background density values (cytoplasm) were subtracted from nuclear and replication foci values, and the ratio of intensity within the viral replication foci relative to the nucleus was calculated. All replication foci assessed (visualized by RPA) were > 0.6 μm in diameter, to distinguish them from small endogenous repair foci often observed in uninfected control cells.

Imaris Colocalization analysis: Z-stack images were first processed in Huygens Essential using the cross-talk corrector and deconvolution with manual background subtraction and default settings. In Imaris, the contour function was used to create a 3D surface of a region of interest drawn around the RPA signal. Each channel was masked based on the 3D surface and utilized for colocalization analysis in Coloc. A threshold was manually determined for each channel and a Coloc channel was generated yielding the Pearson’s coefficient in the ROI volume and the threshold Manders’ coefficient A and B for each set of channel comparisons.

Manual Scoring: To calculate the percentage of HPV replication foci per nuclei containing the protein of interest, replication foci where the protein was visibly enriched relative to the surrounding nucleus were scored and divided by the total number of replication foci observed per nuclei using images collected as a single optical slice. All measured replication foci (visualized by RPA) were over 0.6 μm in size as described above. Using images from maximal projections of 3D optical slices, the number of PML-NBs per nuclei were counted for a minimum of 60 nuclei. The percentage of nuclei where HIRA was associated with PML-NBs was scored visually (for a minimum of 60 nuclei) and classified by the number of PML foci (0 Foci, 1-3 Foci, 4-9 Foci, or 10+ Foci) containing HIRA.

### 5.11 Western blot analysis and antibodies

Prior to keratinocyte extraction, J2-3T3 feeder cells were removed and the keratinocyte monolayer was washed with ice-cold PBS. Cell lysis was performed using 96 ℃ SDS extraction buffer (1% wt/vol SDS, 10 mM Tris-HCl pH 8, 1 mM EDTA pH 8) and protein lysates were sonicated and heat denatured at 95 ℃ for 10 minutes. Protein concentration was determined with a BCA protein assay kit (Thermo -Pierce). 15 µg of protein was supplemented with 50 mM DTT and 1X lithium dodecyl sulfate (LDS) sample buffer (Life Technologies) and heated to 70℃ for 10 minutes. Proteins (12 ug) were separated by electrophoresis on 4-12% Bis-Tris polyacrylamide gels (Invitrogen), transferred to polyvinylidene difluoride (PVDF) membranes and specific proteins detected using the following antibodies: [primary antibodies; Supplemental table 3] HIRA [WC119] mouse monoclonal (Millipore, 04-1488; 1:500 37 ℃ for 1 hour or 4 ℃ overnight), Sp100 rabbit polyclonal (Sigma, HPA016707; 1:2,000), H3.3 rabbit monoclonal (Abcam, ab176840: 1:500 4 ℃ overnight), H3.3S31ph [EPR1873] rabbit monoclonal (Abcam, ab92628-100ul; 1:1,000 4 ℃ overnight), H3 (Millipore, 07-690; 1:5,000 37 ℃ for 1 hour), and GAPDH (6C5) mouse monoclonal (Santa Cruz, sc-32233, 1:1,000 37 ℃ for 1 hour) and [secondary antibodies; goat anti-mouse IgG (H+L), HRP (Invitrogen, 31430; 1:10,000) and goat anti-rabbit IgG (H+L), HRP (Invitrogen, 31460; 1:10,000)]. Proteins were detected with Super Signal West Dura Extended Duration Substrate (Thermo) using a Syngene G:Box. Each membrane was stripped with stripping buffer (GM Biosciences) and reprobed for GAPDH. Protein levels were quantitated using Gene Tools software (version 4.3.9.0) and were normalized to the corresponding value obtained for the GADPH loading control.

## ETHICS STATEMENT

Primary human keratinocytes were isolated from anonymized neonatal foreskins provided to the Dermatology Branch at NIH from local hospitals. The NIH Institutional Review Board (IRB) approved this process and issued an NIH Institutional Review Board waiver.

## ACKNOWLEDGEMENTS

We thank all members of the McBride laboratory for helpful discussions. We acknowledge posthumously the interest and advice on this project by our close colleague Thomas Kristie. This work was supported by the Intramural Research Program of the National Institute of Allergy and Infectious Diseases, of the National Institutes of Health (NIH). The contributions of the NIH authors are considered Works of the United States Government. The findings and conclusions presented in this paper are those of the authors and do not necessarily reflect the views of the NIH or the U.S. Department of Health and Human Services. The majority of this manuscript is derived from the thesis of ADF [71].

