## Supplementary for "Chromatin assembly by the histone chaperone HIRA facilitates Human Papillomavirus replication"

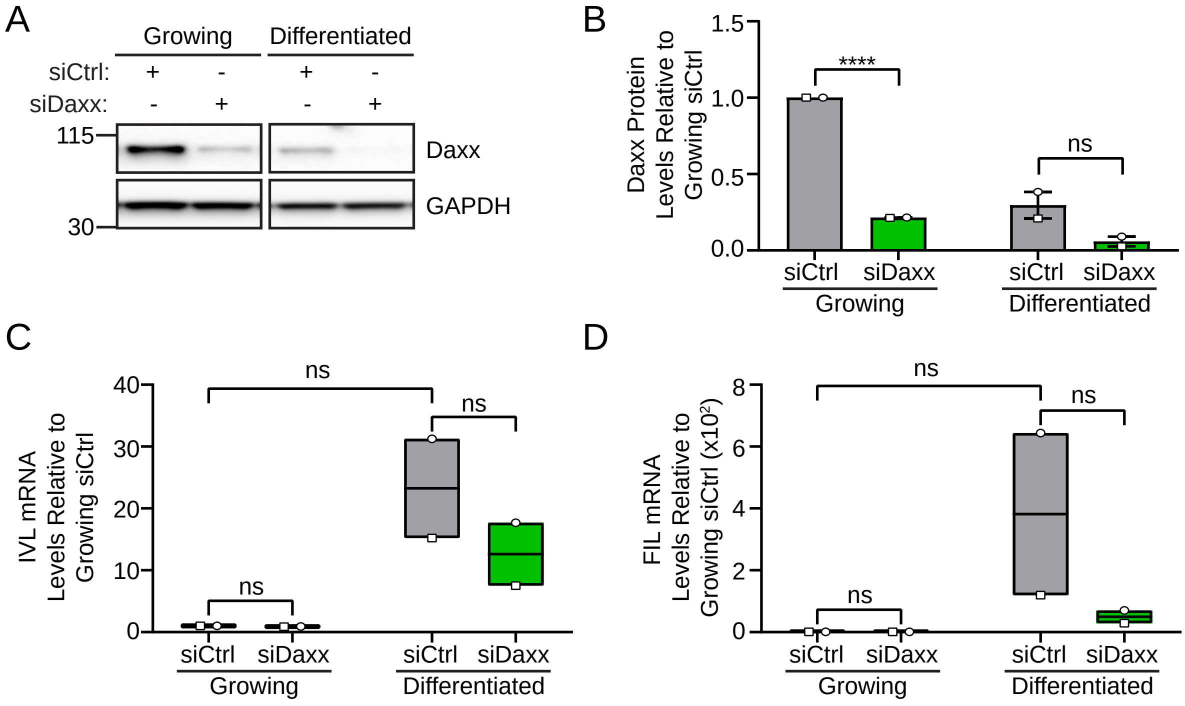

**Supplementary Figure 1: Depletion of Daxx results in impaired keratinocyte differentiation in 9E cells.**

Cells were plated at low density and transfected with 25 nM control (siCtrl) or Daxx (siDaxx) siRNA after 24 hours. Growing and differentiated cells were collected two and eight days post transfection, respectively. **A.** Representative immunoblots of growing and differentiated 9E siCtrl and siDaxx cells, visualizing the protein levels of Daxx and GAPDH (n=2). **B.** Quantitation of panel A, fold change in Daxx levels normalized to GAPDH relative to growing siCtrl (n=2). **C-D.** qRT-PCR using reverse transcribed RNA from growing and differentiated 9E cells transfected with siCtrl or siDaxx, showing the fold change in involucrin and filaggrin cellular transcripts normalized to PPIA within its own condition relative to growing siCtrl (n=2). The same siCtrl samples were utilized in Figure 7. For panel B, error bars represent the range. For panels C-D, floating bars represent minimum and maximum values, and the line indicates the mean. Shapes represent independent experiments. Statistical significance was calculated using an unpaired (Panel B) or paired (Panel C-D) student’s t-test. Ns, not statistically significant, ****, P ≤ 0.0001.

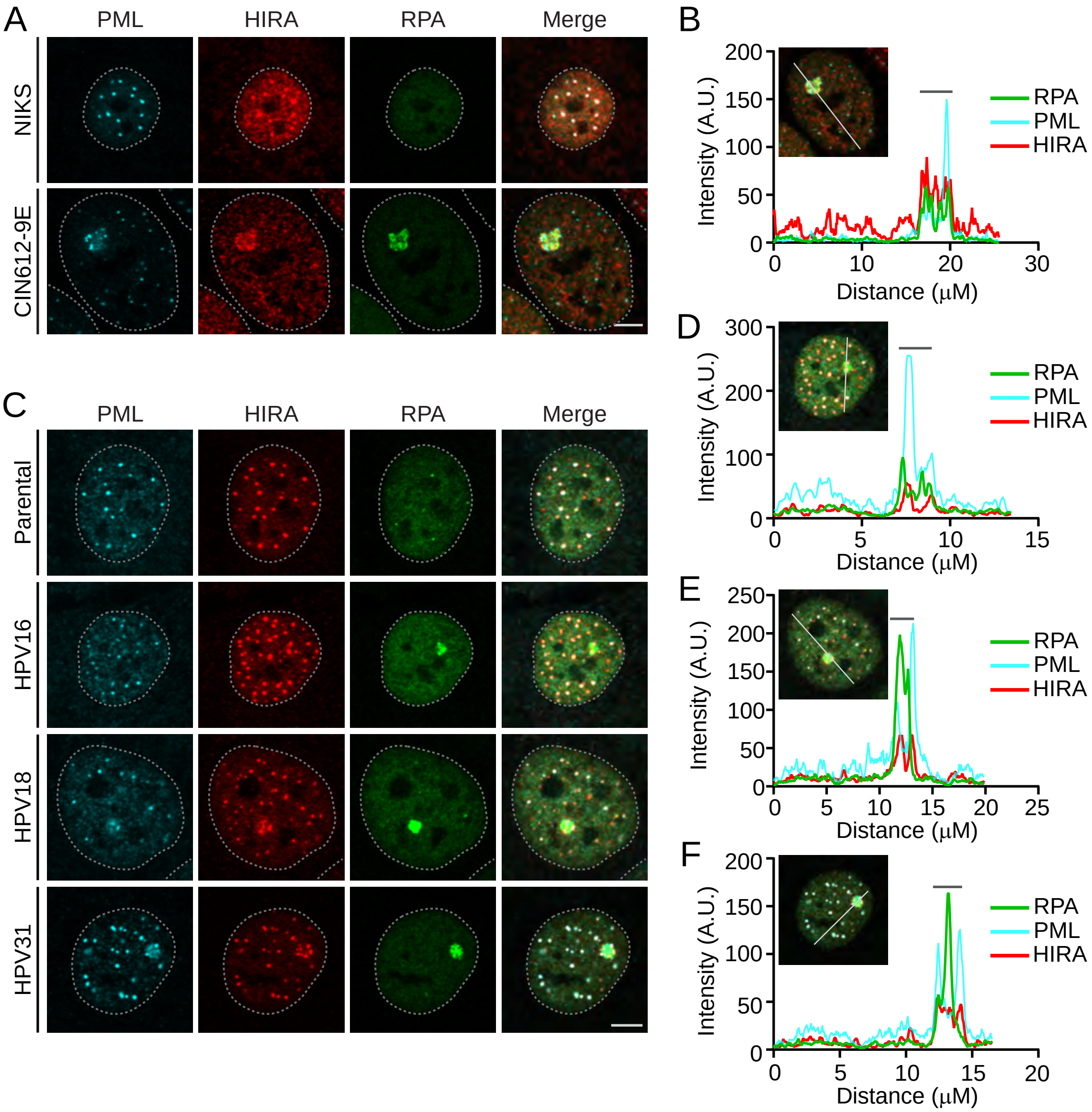

**Supplementary Figure 2: HIRA localizes with PML-NBs and late replication foci in keratinocytes containing various HR-HPV genomes.**

**A.** Representative immunofluorescence staining of PML (pseudo-colored cyan), HIRA (pseudo-colored red), and RPA (green) in differentiated NIKS and 9E cells from three independent experiments. **B.** Fluorescence intensity line scan obtained by drawing a line through the nucleus in panel A in Leica LAS X software. Gray bar above the scan indicates the replication foci. **C.** Representative immunofluorescence staining of PML (pseudo-colored cyan), HIRA (pseudo-colored red), and RPA (green) in differentiated HPV negative HFK, and HFK cells containing HPV16, HPV18, and HPV31 from two independent experiments. **D-F.** Fluorescence intensity line scan obtained by drawing a line through the nucleus in panel C for nuclei containing HPV16, HPV18, and HPV31 genomes, respectively. For all images, a gray dotted line outlines the nucleus defined by DAPI staining. Scale bar, 5 μm.

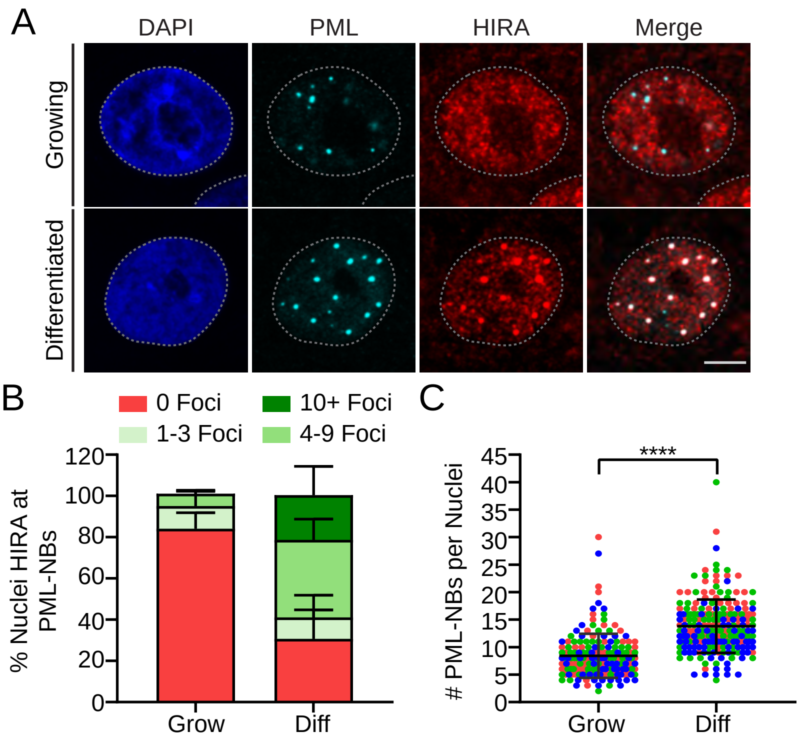

**Supplementary Figure 3: HIRA localization at PML-NBs is increased with differentiation in NIKS cells.**
**A.** Representative immunofluorescence staining of PML (pseudo-colored cyan), HIRA (pseudo-colored red), and RPA (green) in growing and differentiated NIKS cells from three independent experiments. A gray dotted line outlines the nucleus defined by DAPI staining. Scale bar, 5 μm. These images are derived from experiments also represented in Supplementary Figure 1 Panel A. **B.** The percentage of nuclei where HIRA was associated with PML-NBs, categorized by the number of PML-NBs containing HIRA (0 Foci, 1-3 Foci, 4-9 Foci, or 10+ foci). **C.** The number of PML-NBs per nuclei. For panel B and C quantitation, a minimum of 60 nuclei were manually assessed per condition from maximal projections of 3D optical slices. Error bars represent ± standard deviation of the mean and statistical significance was calculated using an unpaired Mann-Whitney test. ****, P ≤ 0.0001.

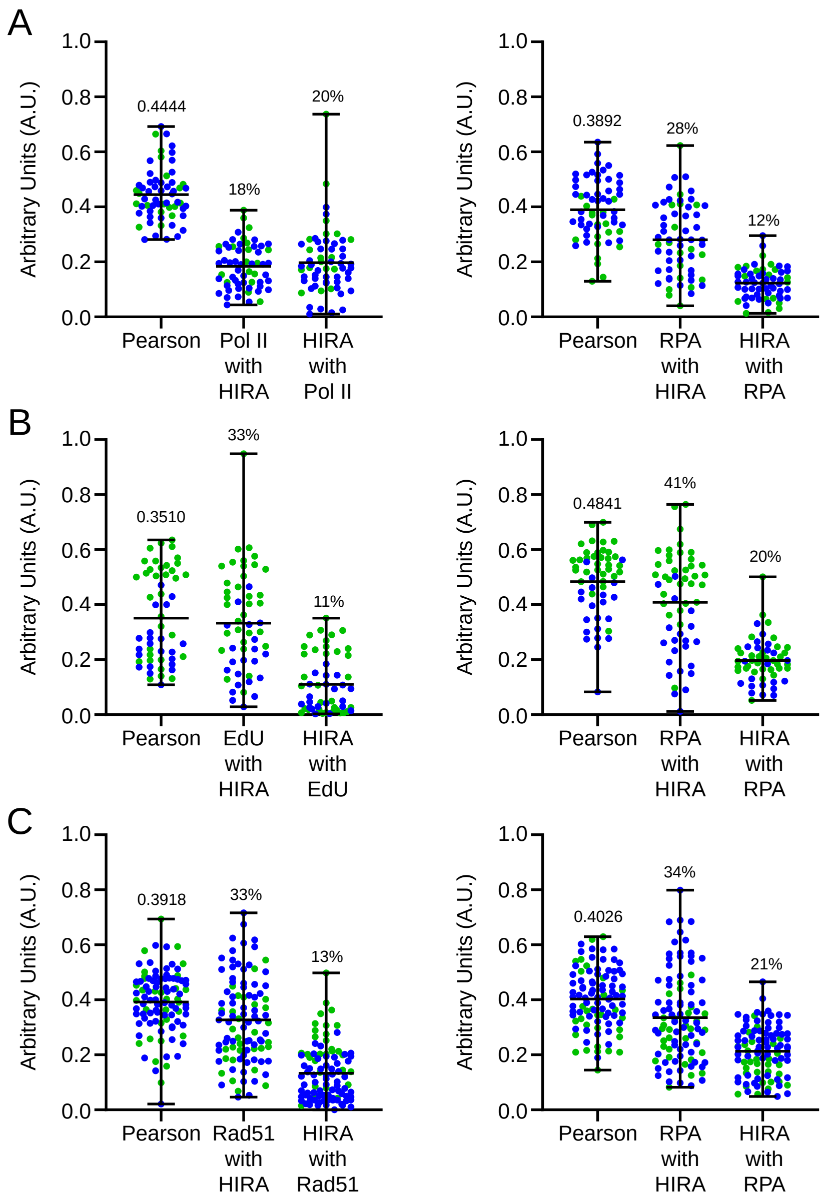

**Supplementary Figure 4: HIRA does not colocalize with any singular host factor at sites of HPV31 replication.**

Z-Stack images were processed in Huygens Essential (cross-talk correction and deconvolution) and colocalization analysis was performed in Imaris within a 3D region of interest (ROI), the replication foci, as defined by the RPA signal. The Pearson's Coefficient in ROI volume and the Threshold Manders' coefficients are shown from images represented in Figure 2. **A.** Colocalization analysis between RNAPIIS2 and HIRA (left) and RPA and HIRA (right) from images represented in Figure 2A-B from a minimum of 11 nuclei/18 replication foci per replicate (n=2). **B.** Colocalization analysis between EdU and HIRA (left) and RPA and HIRA (right) from images represented in Figure 2C-D from a minimum of 12 nuclei/21 replication foci per replicate (n=2). **C.** Colocalization analysis between Rad51 and HIRA (left) and RPA and HIRA (right) from images represented in Figure 2E-F from a minimum of 10 nuclei/30 replication foci per replicate (n=2). Circles denote individual replication foci and colors represent independent experiments. Shown are the mean values and error bars represent the range.

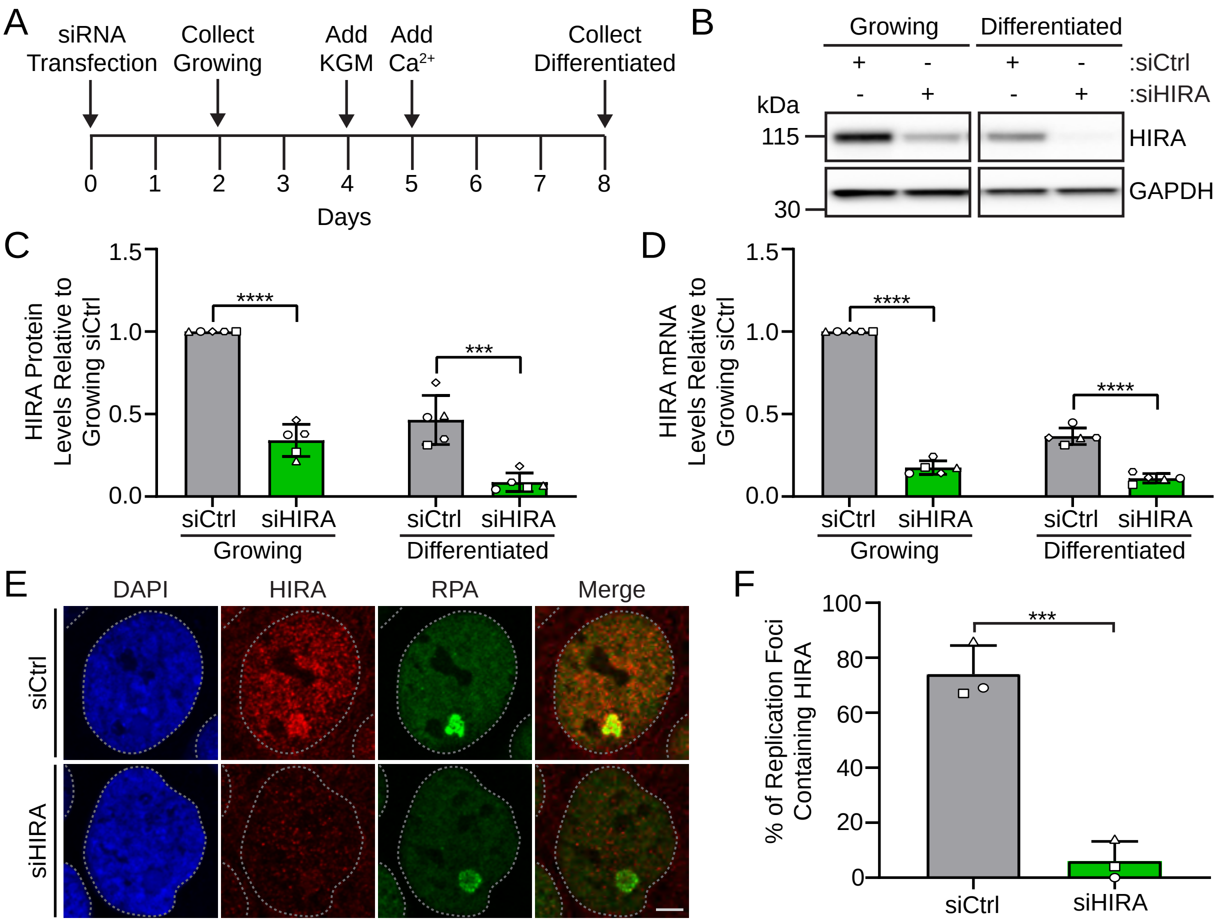

**Supplementary Figure 5: HIRA is efficiently downregulated by siRNA in growing and differentiated 9E cells.**

**A.** Timeline of siRNA knockdown in 9E cells. Cells were plated at low density and transfected with 25 nM control (siCtrl) or HIRA (siHIRA) siRNA after 24 hours. Growing and differentiated cells were collected two, and eight-days post transfection, respectively. **B.** Representative immunoblots of growing and differentiated 9E siCtrl and siHIRA cells, visualizing the protein levels of HIRA and GAPDH (n=5). **C.** Quantitation of panel B, fold change in HIRA levels normalized to GAPDH relative to growing siCtrl (n=5). **D.** qPCR measuring HIRA transcripts, showing the fold change in HIRA mRNA levels relative to growing siCtrl (n=5). **E.** Representative immunofluorescence staining of HIRA (red) and RPA (green) in differentiated 9E cells transfected with siCtrl or siHIRA from three independent experiments. DAPI staining of the nucleus is shown in blue. A gray dotted line outlines the nucleus. Scale bar, 5 μm. **F.** The percentage of replication foci where HIRA is localized to HPV31 replication factories marked by RPA (n = 3; scored minimum of 23 nuclei/30 replication foci). The same siCtrl sample was utilized in Figure 5. Error bars represent ± standard deviation of the mean and shapes represent independent experiments. Statistical significance was calculated using an unpaired student's test. ns, not statistically significant, ***, P ≤ 0.001, **** P ≤ 0.0001.

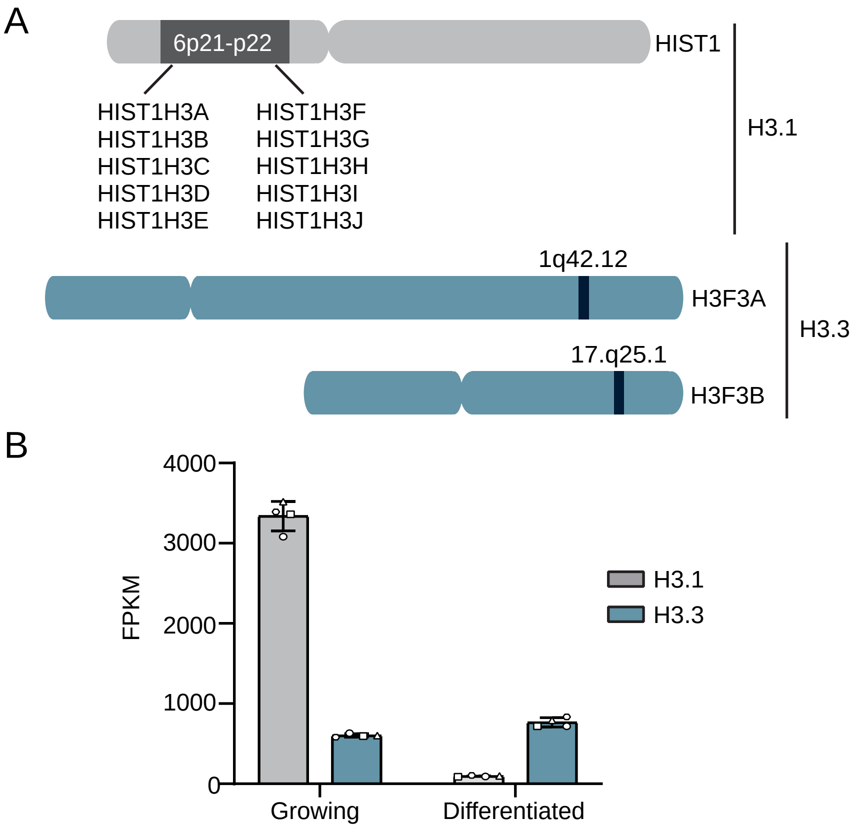

**Supplementary Figure 6: Histone H3.3 is the predominant H3 histone in differentiated keratinocytes.**

**A.** Schematic of the HIST1 gene cluster and H3F3 genes that encode histone H3.1 and H3.3 variants, respectively. The chromosome position of each gene is listed. **B.** RNA-seq analysis was performed on total RNA extracted from growing and differentiated HFK cells containing HPV16, HPV18, and HPV31 genomes described in the methods and the complete dataset will be published elsewhere. Here we show only the combined gene expression (FPKM, Fragments Per Kilobase of transcript per Million mapped reads) of genes encoding histone H3.3 and H3.1 in growing and differentiated HFKs (n=4 technical replicates). Error bars represent ± standard deviation of the mean and shapes represent independent replicate values.

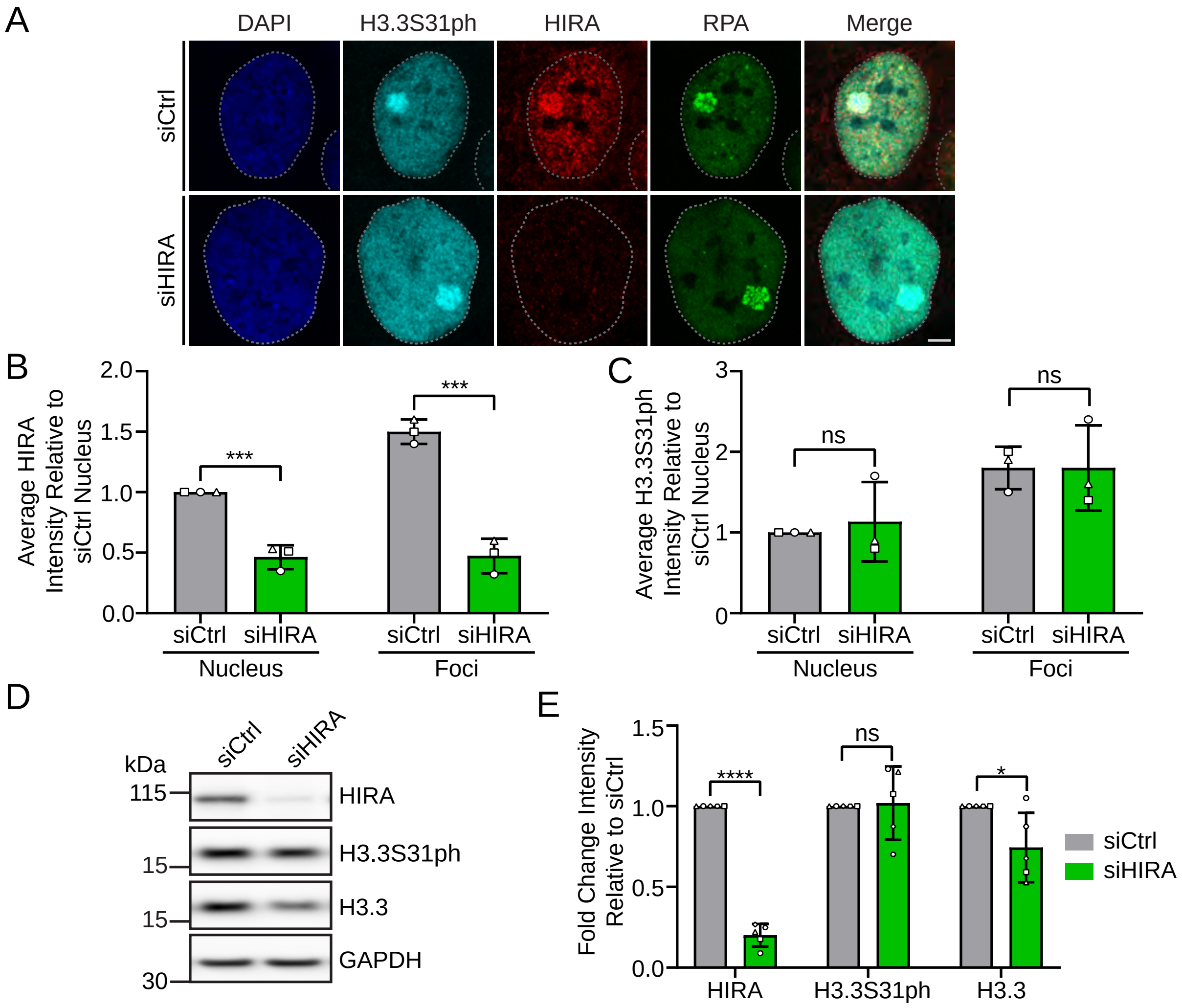

**Supplementary Figure 7: Enrichment of histone variant H3.3 phosphorylated at serine 31 at HPV31 replication foci is independent of HIRA.**

**A.** Representative immunofluorescence staining of H3.3S31ph (cyan), HIRA (red), and RPA (green) in differentiated siCtrl and siHIRA 9E cells from three independent experiments. DAPI staining of the nucleus is shown in blue. A gray dotted line outlines the nucleus. Scale bar, 5 μm. Average HIRA (**Panel B**) and H3.3S31ph (**Panel C**) intensity at RPA foci and in the nucleus relative to siCtrl nucleus was measured in ImageJ. **D.** Representative immunoblots of differentiated 9E siCtrl and siHIRA cells, visualizing the protein levels of HIRA, H3.3S31ph, H3.3, and GAPDH (n = 5). The GAPDH blot shown is representative of all blots but matched specifically to that of the H3.3 blot. **E.** Quantitation of panel D, showing the fold change in protein levels normalized to GAPDH relative to siCtrl (n= 5). Quantitation of the HIRA westerns are also represented in Supplementary Figure 4. For panel B and C quantitation, n=3; measured a minimum of 31 replication foci/18 nuclei per experiment. Error bars represent ± standard error of the mean (intensity quantitation) or standard deviation of the mean (western blot quantitation) and shapes represent independent experiments. Statistical significance was calculated an unpaired student's t-test. ns, not statistically significant, *, P ≤ 0.05, ***, P ≤ 0.001, ****, P ≤ 0.0001.

**Supplementary Table 1: siRNA sequences used in this study**

| Target mRNA | Sequence (5' - 3') |
|---|---|
| Non-targeting Control | UGGUUUACAUGUCGACUAA |
| | UGGUUUACAUGUUGUGUGA |
| | UGGUUUACAUGUUUUCUGA |
| | UGGUUUACAUGUUUUCCUA |
| Sp100 | GAAGUGAGCCUGUGAUCAA |
| | AGGCAUAGAUCUAAAGUAA |
| | UACCAGAGCCCAUGGAUUU |
| | GAAGGGCACUCUAUAUAAG |
| HIRA | CAUGGGACCCUGUUGGUAA |
| | GGAUAACACUGUCGUCAUC |
| | GCUCCGAUCCUUCCAUGUA |
| | GCAGGCGAUUCUGUCAAUA |

**Supplementary Table 2: Primers**

| Target | Nucleotide Position | Strand | Sequence (5' - 3') | Reference |
|---|---|---|---|---|
| Detection of HPV31 9E genome with qPCR | 618-640 | Sense | CTGACCTCCACTGTTATGAGCAA | [33] |
| Detection of HPV31 9E genome with qPCR | 686-663 | Anti-sense | CAGCTGGACTGTCTATGACATCCT | [33] |
| qPCR primer for (H1RNA) gene (RPPH1) on chr 14, cytoband 14q11.2. | 7-26 | Sense | CGGAGGGAAGCTCATCAGTG | |
| qPCR primer for (H1RNA) gene (RPPH1) on chr 14, cytoband 14q11.2. | 95-76 | Anti-sense | TGGCCCTAGTCTCAGACCTT | |
| qPCR primer for HPV31 E6*1 | 186-210, 413-416 | Sense | AGATTGAATTGTGTCTACTGCAAAGGTGT | [33] |
| qPCR primer for HPV31 E6*1 | 520-498 | Anti-sense | GCTATGCAACGTCCTGTCCACCT | [33] |
| qPCR primer for HPV31 E1^E4 | 857-877, 3294-3296 | Sense | CTACAATGGCTGATCCAGCAGCA | [33] |
| qPCR primer for HPV31 E1^E4 | 3405-3387 | Anti-sense | CGCCGCACACCTTCACTGG | [33] |
| qPCR primer for HPV31 L1 | 3562-3590, 5552-5554 | Sense | TGCAACTACACCTATAATACACTTAAAAGATG | [33] |
| qPCR primer for HPV31 L1 | 5641-5619 | Anti-sense | TCGTGTTACATATTCATCCGTGC | [33] |
| qPCR primer for Human involucrin in differentiating cells | 31-53 | Sense | TTACTGTGAGTCTGGTTGACAGT | |
| qPCR primer for Human involucrin in differentiating cells | 150-128 | Anti-sense | GGTATTGACTGGAGGAGGAACAG | |
| qPCR primer for Human filaggrin in differentiating cells | 198-221 | Sense | GCAAATCCTGAAGAATCCAGATGA | |
| qPCR primer for Human filaggrin in differentiating cells | 324-303 | Anti-sense | TGCTTGAGCCAACTTGAATACC | |
| qPCR primer for HIRA | 280-299 | Sense | GGGTCAACCACAATGGCAAG | |
| qPCR primer for HIRA | 351-332 | Anti-sense | CTCCAGTTGCGAACTTGGTC | |
| qPCR primer for Sp100, Isoform A | 1594-1613 | Sense | ACTTGGCCTGCAGAATGTCA | [33] |
| qPCR primer for Sp100, Isoform A | 1676-1655 | Anti-sense | CAAGGTAGTGAAGGTGCTCAGA | [33] |
| qPCR primer for Sp100, Isoform B | 2212-2236 | Sense | TCTGCCAATGTCTCGTCTATTATGT | [33] |
| qPCR primer for Sp100, Isoform B | 2291-2263 | Anti-sense | TTATGATGATGGGTCAATTTAAAGACTGT | [33] |
| qPCR primer for Sp100, Isoform C | 2455-2472 | Sense | CTGCCTGAGGAGCAGTTGAA | [33] |
| qPCR primer for Sp100, Isoform C | 2538-2519 | Anti-sense | CGGTTCTGAGGCGAAAAAGC | [33] |
| qPCR primer for Sp100, Isoform HMG | 2224-2244 | Sense | GTTGACCCTTGTGAGGAGCAT | [33] |
| qPCR primer for Sp100, Isoform HMG | 2365-2345 | Anti-sense | TGTCCGCCTTTGCCATATCTT | [33] |
| qPCR primer for cyclophilin PPIA | 503-526 | Sense | AGAACTTCATCCTAAAGCATACGG | |

| | | | |
|---|---|---|---|
| qPCR primer for cyclophilin PPIA | 532-513 | Anti-sense | TGCTTGCCATCCAACCACTC |

**Supplementary Table 3: Antibodies**

| Target Protein | Manufacturer | Catalog Number | RRID Number | Species | IF Concentration | Western Blot Concentration | Western Blot Conditions |
|---|---|---|---|---|---|---|---|
| HIRA (WC119)* | Millipore | 04-1488 | AB_1977097 | Mouse Monoclonal | 1:100 | 1:500 | 5% Milk in TBST; 4C overnight |
| PML (H-238) | Santa Cruz | sc-5621 | AB_2166848 | Rabbit Polyclonal | 1:100 | | |
| Sp100 | Sigma | HPA016707 | AB_1857398 | Rabbit Polyclonal | 1:500 | 1:2,000 | 5% Milk in TBST; RT 1 hour |
| UBN1 | Sigma | HPA061029 | AB_2684420 | Rabbit Polyclonal | 1:100 | | |
| ASF1a | Cell Signaling | 2990s | AB_2289918 | Rabbit Monoclonal | 1:100 | | |
| RNA Pol II S2 | Abcam | ab-5095 | AB_304749 | Rabbit Polyclonal | 1:100 | | |
| Rad51 | Abcam | ab133534 | AB_2722613 | Rabbit Monoclonal | 1:100 | | |
| H3.3S31ph (EPR1873) | Abcam | ab92628-1001 | AB_10563637 | Rabbit Monoclonal | 1:100 | 1:1,000 | 5% BSA in TBST; 4C overnight |
| H3.3 | Abcam | ab176840 | AB_2715502 | Rabbit Monoclonal | 1:200 | 1:500 | 5% Milk in TBST; 4C overnight |
| H3 | Millipore Sigma | 07-690 | AB_417398 | Rabbit Polyclonal | | 1:5,000 | 5% Milk in TBST; RT 1 hour |
| RPA32/RPA2 (4E4) | Cell Signaling | 2208 | AB_2927646 | Rat Monoclonal | 1:300 | | |
| Glu-Glu Epitope Tag | Novus | NB600-354 | AB_10003158 | Rabbit Polyclonal | 1:300 | | |
| GAPDH (6C5) | Santa Cruz | sc-32233 | AB_627679 | Mouse Monoclonal | | 1:1,000 | 5% Milk in TBST; RT 1 hour |
| Alexa Fluor 488 H2AX (pS139) | BD Pharmingen | 560445 | AB_1645352 | Mouse | 1:10 | | |
| TopBP1 (R1180) | Boner et. al 2002 | | | Rabbit | 1:500 | | |
| TopBP1 (B-7) | Santa Cruz | sc-2710443 | AB_10610636 | Mouse Monoclonal | 1:100 | | |
| ATR | Cell Signaling | 2790 | AB_2227860 | Rabbit Polyclonal | 1:100 | | |
| Phospho-Chk1 (Ser345) (133D3) | Cell Signaling | 2348 | AB_331212 | Rabbit Monoclonal | 1:200 | | |

| | | | | | |
|---|---|---|---|---|---|
| H3K27ac | Sigma | 07-360 | AB_310550 | Rabbit Polyclonal | 1:100 |
| p300 (3G230/NM-11) | Abcam | ab14984 | AB_301550 | Mouse Monoclonal | 1:500 |
| Brd4 (1F11) | Cheng-Ming Chiang | | | Mouse Monoclonal | 1:100 |

* Antibody incubated at room temperature for 1 hour or at 4C overnight.
